# Structural basis of stepwise G protein activation by the viral chemokine receptor US28

**DOI:** 10.64898/2025.12.10.692789

**Authors:** Kevin M. Jude, Carl-Mikael Suomivuori, Deepa Waghray, Shoji Maeda, Yoshinori Fujiyoshi, Asuka Inoue, K. Christopher Garcia, Naotaka Tsutsumi

**Author notes:** Correspondence (N.T.), (K.C.G.), (A.I.), (C.-M.S.).

## Abstract

G protein-coupled receptors (GPCRs) govern diverse cellular responses and are crucial drug targets. However, the sequence of structural events from G protein recognition to GDP release has remained elusive. Here, we leveraged the viral chemokine GPCR US28 to capture multiple activation states of the US28-G_q_ complex. Using cryo-electron microscopy and an engineered chemokine superagonist, we determined three distinct complex structures, capturing a GDP-bound Encounter state, the nucleotide-free Canonical state, and a putative intermediate bridging the two states, the Transition-to-Canonical state. These structures, along with simulations and functional data, provide high-resolution snapshots of a plausible G protein activation trajectory and support a stepwise conformational model for G protein activation. This activation cascade closely parallels mechanisms proposed for human GPCRs, suggesting a conserved GPCR signaling mechanism.

## Introduction

G protein-coupled receptors (GPCRs) constitute the largest family of integral membrane proteins in the human genome and are responsible for transducing an immense variety of extracellular stimuli, including photons, odors, tastes, hormones, and neurotransmitters, into intracellular signaling^1^. Thus, G protein signaling underpins nearly all aspects of human physiology, from sensory perception to cardiovascular regulation and immune function. Consequently, GPCRs represent the targets for approximately one-third of all approved drugs, making the elucidation of their activation mechanisms a cornerstone of modern medicine and pharmacology^2,3^.

GPCRs function as guanine nucleotide exchange factors (GEFs) for their cognate heterotrimeric G proteins, which are composed of Gα and Gβγ subunits^4,5^. In the basal state, the G protein is inactive with GDP bound to the Gα subunit. Upon activation by an extracellular stimulus, the GPCR engages the G protein and induces a conformational change in the Gα subunit that weakens the affinity for GDP. The subsequent GDP release and rapid binding of the more abundant intracellular GTP lead to the dissociation of the Gα and Gβγ subunits^4,6^, which then modulate downstream effectors to elicit a specific cellular response^7,8^.

The recent revolution in cryo-electron microscopy (cryo-EM) has led to a surge in high-resolution structures of GPCRs coupled to G proteins^9^. However, this wealth of structural information is heavily biased toward a specific endpoint in the activation sequence: the fully engaged, nucleotide-free complex^10^. This state, representing the complex after GDP release but prior to GTP binding, corresponds to an energetic minimum *in vitro* and is thus more amenable to structural visualization, although it is transient in a cellular environment^11^. As a result, the initial recognition of the inactive, GDP-bound G protein by the receptor and the subsequent conformational transitions that catalyze GDP release have remained structurally elusive^12,13^. Insights into these early events have thus far been primarily derived from molecular dynamics (MD) simulations^14–16^. Although recent studies have visualized later events in the G protein cycle, such as post-GTP-binding rearrangements^11,17^, a comprehensive structural understanding still requires visualization of the full activation trajectory, beginning with the initial molecular engagement between GPCR and G protein.

To bridge this mechanistic gap, we turned to US28, a human cytomegalovirus (HCMV)-encoded GPCR. US28 is a viral chemokine receptor known to promiscuously engage host proteins to subvert cellular signaling for the benefit of the HCMV lifecycle^18–22^. It binds a wide array of extracellular CC and CX_3_C chemokines and interacts with multiple intracellular G protein families, including G_q/11_, G_12/13_, and G_i/o_^21^. We previously determined crystal structures of US28 in its ligand-free state and bound to the native or engineered variants of the CX_3_CL1 chemokine^23,24^. We also reported cryo-EM structures of CX_3_CL1-US28 complexed with G_i_ or G_11_^13,22^. Strikingly, one of these structures captured a state where GDP remains bound to Gα_11_ in its US28-G_11_ complex^13^. We originally termed this the transition-like state (TL-state), as it likely represents an initial engagement preceding nucleotide release. This finding highlighted US28 as a powerful model for visualizing metastable states of the GPCR-G protein activation cycle beyond the canonical nucleotide-free complex structure. However, the corresponding ‘canonical state’ (C-state) for the US28-G_q/11_ complex—which represents the terminal step of GDP release before GTP binding—had not been elucidated, precluding a direct, side-by-side structural comparison across the entire activation process.

In this work, we present the cryo-EM structures of the US28-G_q_ complex activated by the engineered chemokine superagonist, CX_3_CL1.44^24^, unveiling several key snapshots during the stepwise G_q_ activation (Fig. 1). We report not only the nucleotide-free C-state but also the previously reported GDP-bound state at higher resolution, which we redefine as the Encounter state (E-state) hereafter. Furthermore, we describe a putative structural intermediate between these two, which we designate the ‘Transition-to-Canonical’ state (T2C-state). Based on comparative analysis, we propose that the transition from initial engagement to the nucleotide-free state is driven by α5 engagement, which acts through an allosteric conduit to disrupt the salt bridge between the conserved glutamate and arginine in the P-loop and Switch I, respectively. This downstream disruption initiates a salt bridge relay that destabilizes the inter-domain interface. The higher-resolution E-state clarifies the initial receptor-G protein encounter, while the C-state complex elucidates the structural basis for US28’s G protein-subtype selectivity. MD simulations and structure-guided mutagenesis provide complementary data consistent with the relevance of these conformations during G protein activation process. Finally, the proposed mechanism closely resembles that suggested for G_s_ proteins catalyzed by human GPCRs^16,25^, supporting a conserved mechanism of nucleotide exchange across GPCR families. Collectively, these findings establish an integrated structural framework for understanding the sequence of molecular events that drive GPCR-mediated nucleotide exchange.

**Fig. 1.**
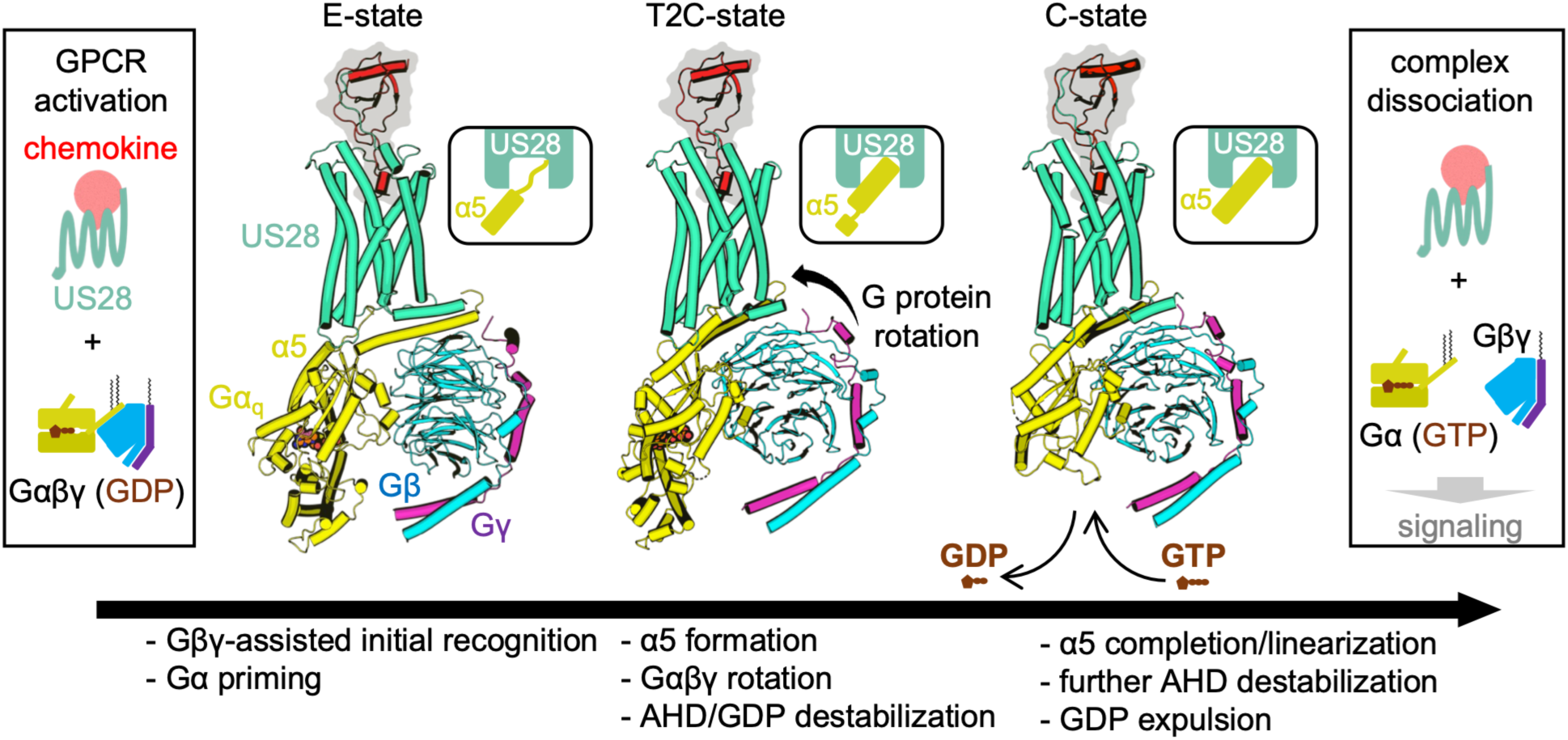
Model of the stepwise G_q_ activation by US28. Schematic representation of the proposed G protein activation trajectory. The structures are colored as follows: US28 (green-cyan), chemokine (red with gray surface background), Gα_q_ (yellow), Gβ (cyan), and Gγ (purple). Upon GPCR (US28) activation, the inactive, GDP-bound Gαβγ heterotrimer is recruited. The complex transitions through the E-state, the T2C-state, and the C-state. The E-state represents the initial engagement where the receptor bridges the Gα and Gβγ subunits, characterized by a loosely anchored G protein with a non-helical Gα_q_ C-terminus. During the transition from the E-state to the T2C-state, the G protein undergoes a rotational movement relative to the receptor as the α5 helix forms and inserts into the receptor core, although its base remains broken. Subsequently, in the C-state, the α5 helix is completed, leading to large-scale Gα domain separation and GDP release. GTP binding to the C-state leads to complex dissociation and downstream signaling. The insets highlight the stepwise conformational remodeling of the Gα_q_ C-terminal α5-helix (yellow) and its engagement with the US28 intracellular core (green-cyan). Key molecular events, such as α5 formation, G protein rotation, and GDP expulsion, are summarized at the bottom.

## Results

### Overall structure of the active US28-G_q_ complex at multiple states

To obtain US28-G_q_ complexes in both GDP-bound and nucleotide-free states from a single sample, we used the G protein-biased chemokine variant CX_3_CL1.44^13,24^. The complex was formed in detergent micelles, a condition known to yield the nucleotide-free CX_3_CL1.44-US28-G_11_ complex and to allow GTP turnover catalyzed by US28^13^. This micellar environment is also commonly used for “time-resolved” experiments to track nucleotide-induced conformational changes, as it slows GDP release compared to a lipid bilayer^26^. We also used a chimeric Gα_q_ (Gα_qiN18_), in which the first 18 N-terminal residues are replaced by the corresponding Gα_i1_ sequence, to enable the binding of the scFv16 fiducial marker^27^, yielding a viable sample for cryo-EM imaging (Supplementary Fig. 1a,b).

Our initial cryo-EM analysis revealed that the sample contained both “monomeric” and “dimeric” forms of the US28-G_q_ complex (Supplementary Fig. 1c,d). Focusing on the monomeric particles, we performed extensive classification prioritized on the US28-G_q_ interface, yielding a series of well-resolved 3D reconstructions (Supplementary Fig. 2-4). Within this “monomeric” population, we resolved at least three distinct states that differed principally in the conformation of the Gα_q_ subunit and the presence or absence of GDP (Fig. 2a,b). Despite this variability in the G protein, US28 maintained a highly consistent active seven-transmembrane conformation across all states, with Cα root mean square deviations (RMSDs) ranging from 0.58 to 0.63 Å among the 227 transmembrane α-helical residues of the receptor. Importantly, we observed no macroscopic structural rearrangements among these states (Supplementary Fig. 5a). This structural conservation suggests that fully activated US28 functions primarily as a rigid catalytic scaffold throughout the progressive stages of G protein engagement.

**Fig. 2.**
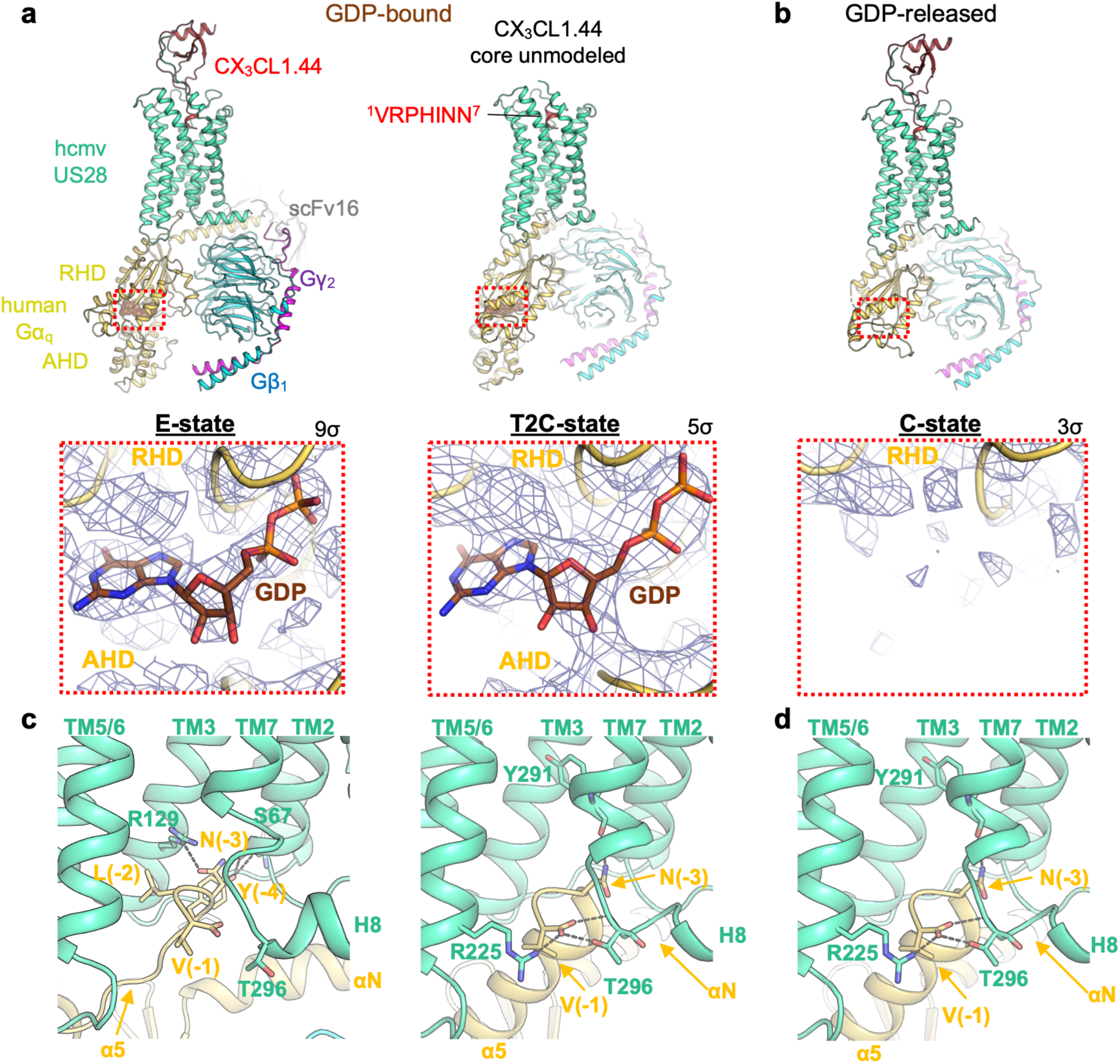
Structural snapshots of the plausible US28-G_q_ activation trajectory. **a**, Structures of the CX_3_CL1.44-US28-G_q_ complex in GDP-bound states, the E-state and the T2C-state. **b**, Structure of the nucleotide-free state, the C-state. Color scheme is as follows: US28 (green-cyan), CX_3_CL1.44 (red), Gα_q_ (yellow), Gβ1 (cyan), Gγ2 (purple), and scFv16 (gray). Bottom panels show close-up views of the nucleotide-binding pocket with cryo-EM density (blue mesh) with the GDP model shown as sticks and surrounding region shown as ribbon diagram. Map contour levels (σ) are indicated in the figure. **c, d,** Close-up views of the interface between the US28 intracellular core (green-cyan) and the Gα_q_ C-terminus (yellow) in **c**, the E- and T2C-states, and **d**, the C-state. Key interacting residues are shown as sticks. Polar interactions (hydrogen bonds or salt bridges) are indicated by dashed lines.

The first prominent state, which we termed the E-state, represents a major GDP-bound conformation^13^. Here, the G_q_ protein retains GDP within a pocket enclosed by its Ras-homology domain (RHD) and α-helical domain (AHD) while initiating engagement with the receptor (Fig. 2a,c). This structure, refined to 2.6 Å resolution, is analogous to the previously reported TL-state CX_3_CL1-US28-G_11_ structure but at substantially higher resolution^13^, thereby enabling more accurate modeling. A key feature is that the Gα_q_ α5 C-terminus adopts a non-helical yet partially ordered conformation and is loosely anchored to the US28 intracellular core, rather than forming the canonical α-helix that mediates extensive contacts in the fully active state. US28 bridges the Gα_q_ and Gβ subunits through multi-point interactions, including contacts between US28 intracellular loop (ICL) 2 and the Gα_q_-αN helix, as well as between US28-ICL1/helix 8 (H8) and Gβ (Supplementary Fig. 5b). This bridging architecture likely represents the Gβ-assisted, high-affinity initial engagement of the GPCR-G protein complex.

Concurrently, we resolved a set of nucleotide-free conformations, which are collectively referred to as the canonical state (C-state), based on their close resemblance to the majority of active GPCR-G protein complex structures^11,13^. The C-state is characterized by closer packing between US28 and the Gα_q_-RHD, the formation of the canonical α5 helix at the Gα_q_ C-terminus, and the separation of the AHD from the RHD (Fig. 2b,d). While this C-state population exhibited minor stochastic heterogeneity, we focused on the particle subset that yielded the highest-quality map at 2.6 Å resolution (Supplementary Fig. 2).

Notably, 3D classification resolved a putative intermediate bridging these two states: the “Transition-to-Canonical” state (T2C-state), refined to 2.9 Å resolution (Fig. 2a). This state exhibits hybrid features: while the C-terminus of Gα_q_-α5 forms an α-helix resembling the C-state (Fig. 2c right and Fig. 2d), the protein still retains GDP within the AHD-RHD interface, characteristic of the E-state (Fig. 2a bottom). Furthermore, the membrane-distal (N-terminal) segment of the α5 helix is broken, a conformational feature that distinguishes the T2C-state from the other states (Fig. 1 cartoon insets) and that is discussed in more detail below. These features indicate that the T2C-state represents a plausible on-pathway intermediate bridging the E-state and C-state.

Consistent with the observed stochastic flexibility between US28 and G_q_, as well as between Gα_q_-RHD and G_q_-AHD, we also obtained additional 3D reconstructions related to the C- state (Supplementary Fig. 2 and 5b). These include variations of the C-state and a distinct nucleotide-free state with a closed AHD (C’-state), which are discussed in the Supplementary Discussion.

### MD simulations support the T2C-state as a plausible on-pathway intermediate

To investigate the conformational dynamics of the T2C-state and its structural relationship to the E- and C-states, we turned to all-atom MD simulations (Fig. 3a-c and Supplementary Fig. 6). We performed 16 simulations, each around 2 µs in length, initiated from the T2C-state embedded in a lipid bilayer (see Methods, Supplementary Tables 2 and Supplementary Fig. 6).

**Fig. 3.**
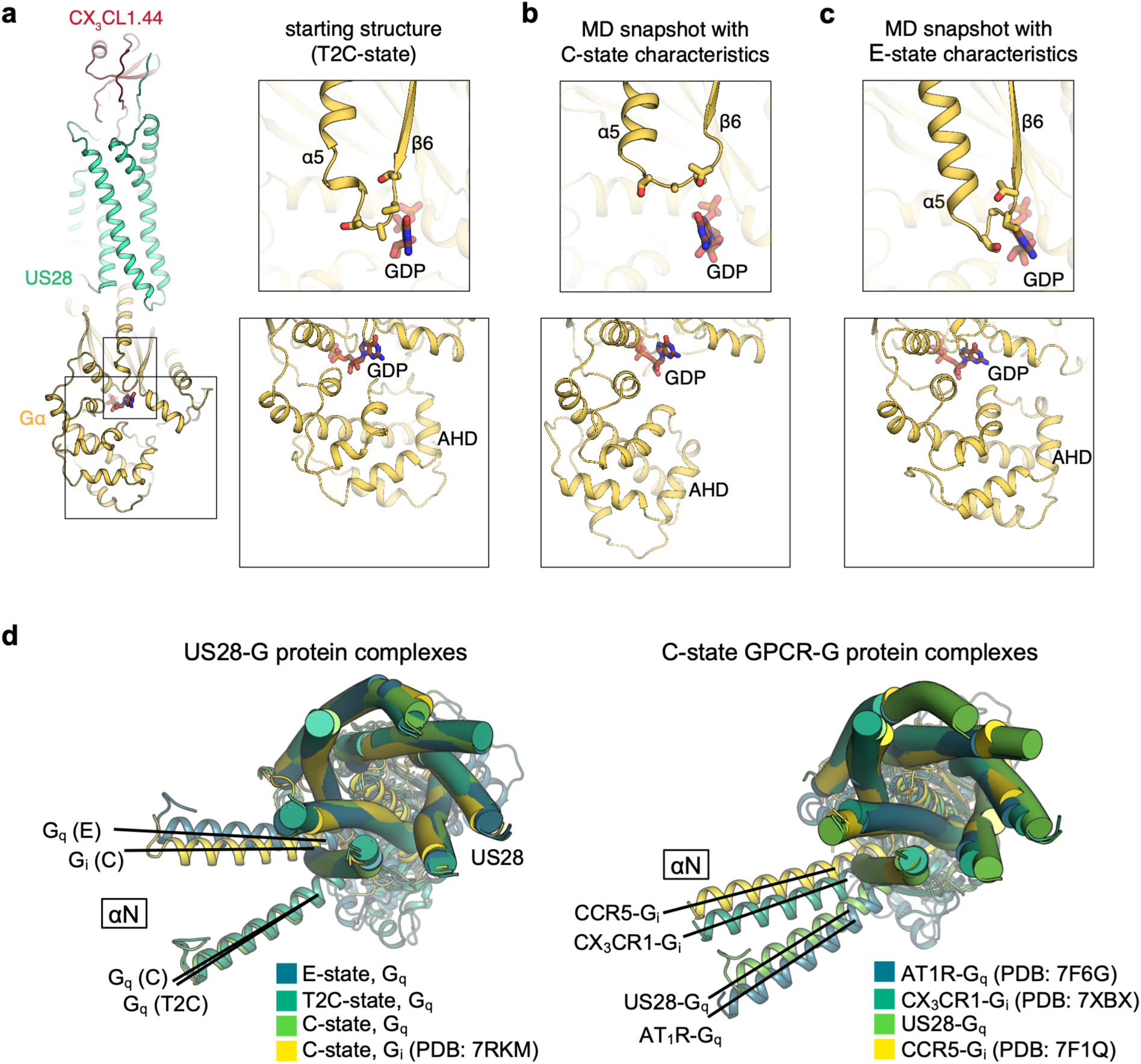
Dynamics of the T2C intermediate and G protein rotation. **a-c**, Simulations reveal partial transitions from the T2C-state toward the C- and E-states. **a**, The T2C-state structure used as a starting point for MD simulations. The T2C-state is characterized by, e.g., a broken membrane-distal segment of α5, with the TCAT motif (shown in sticks) of the β6-α5 loop in close proximity to GDP (top), as well as by the AHD being in close proximity to the RHD (bottom). Gβ and Gγ are omitted for clarity. **b**, Representative frame of a simulation with a partial transition from the T2C-state toward the C-state. In this simulation, the membrane-distal segment of α5 regains helicity in a way that moves the TCAT motif in the β6-α5 loop away from GDP, and the AHD separates from the RHD. This frame is taken from *t* = 0.8 µs of simulation 16. **c**, Representative frame of a simulation with a partial transition from the T2C-state toward the E-state. In this simulation, the membrane-distal segment of α5 regains helicity in a way that keeps the TCAT motif close to GDP, and the AHD remains in proximity to the RHD. This frame is taken from *t* = 0.8 µs of simulation 4. For time traces of relevant metrics for all simulations, see Supplementary Fig. 6. **d**, Overall conformational rearrangements during G_q_ activation and comparison with other GPCRs. Left: Superposition of the US28-Gα_q_ complexes (E-state, T2C-state, and C-state) and the US28-Gα_i_ complex (C-state, PDB: 7RKM), aligned on the US28 region (E-state shown), showing the rotation of the Gα protein during the activation trajectory. Right: Comparison of the C-state US28-Gα_q_ complex with representative C-state GPCR-Gα protein complexes (AT1R-Gα_q_, PDB: 7F6G; CX_3_CR1-Gα_i_, PDB: 7XBX; CCR5-Gα_i_, PDB: 7F1Q), highlighting variations in Gα protein engagement mode. Only aligned TM helices of US28 are shown, as cylinders, for clarity.

In 6 simulations, we identified partial transitions toward the C-state. While the membrane-distal segment of α5 is broken in the starting T2C-state structure (Fig. 3a), we observed this segment regaining helicity such that the TCAT motif in the β6-α5 loop moves away from the nucleotide-binding site, which is characteristic of the C-state (Fig. 3b and Supplementary Fig. 6a). Furthermore, in 1 out of those 6 simulations, we additionally observed large-scale separation of the AHD from the RHD and subsequent AHD mobility, also characteristic of the C-state (Fig. 3b and Supplementary Fig. 6b). We did not observe GDP release at the simulated timescales.

In addition to these 6 simulations, we also identified what we consider a partial transition toward the E-state in a single simulation (Fig. 3c and Supplementary Fig. 6). In this simulation, the membrane-distal segment of α5 also regains helicity, but in a way that keeps the TCAT motif close to GDP, in a manner resembling the E-state (Supplementary Fig. 6c). However, we note that the α5 C-terminal segment remains helical in this simulation, distinguishing it from the unwound conformation observed in the experimental E-state structure.

The remaining 9 simulations generally retained characteristics of the T2C-state, including a broken membrane-distal segment of α5 and the AHD remaining in proximity to the RHD (Supplementary Fig. 6). The observation that the T2C-state was stably maintained in a lipid bilayer over these timescales supports that it represents a physically viable conformation rather than a micelle-induced artifact.

These bidirectional transitions from the T2C-state toward both the E- and C-states observed in simulations are consistent with the T2C-state representing a plausible on-pathway intermediate between the E- and C-states.

### Stepwise rearrangement of the receptor-G protein interface

A substantial conformational rearrangement at the US28-G_q_ interface distinguishes the E-state from the T2C- and C-states. This rearrangement is primarily driven by the formation of the α5 helix within Gα_q_ and corresponding shifts in the docking mode. When viewed along the CX_3_CL1.44-US28-Gα_q_ axis, G_q_ rotates by approximately 33^°^ accompanied by a slight translation of less than 2 Å along the αN helix (Fig. 3d left), a motion analogous to the GTP-induced dissociation of the β2-adrenergic receptor (β2AR)-G_s_ complex^11^. These G protein orientations fall within the range observed for similar class A GPCR complexes^28–30^ (Fig. 3d right). As a result, the interface areas and the residue-level contacts are altered. US28’s buried surface areas for the Gα_q_/Gβ_1_ interfaces are 1206/191 Å^2^, 1124/39 Å^2^, and 1031/56 Å^2^ for the E-state, T2C-state, and C-state, respectively. Interestingly, this rotation, coupled with conformational changes in Gα_q_, results in a rearrangement of the interacting residue pairs and the overall interface, primarily at Gα_q_-αN and Gα_q_-α5, even though the fundamental set of Gα_q_-binding residues on US28 remains largely consistent (Fig. 4a and Supplementary Fig. 7).

**Fig. 4.**
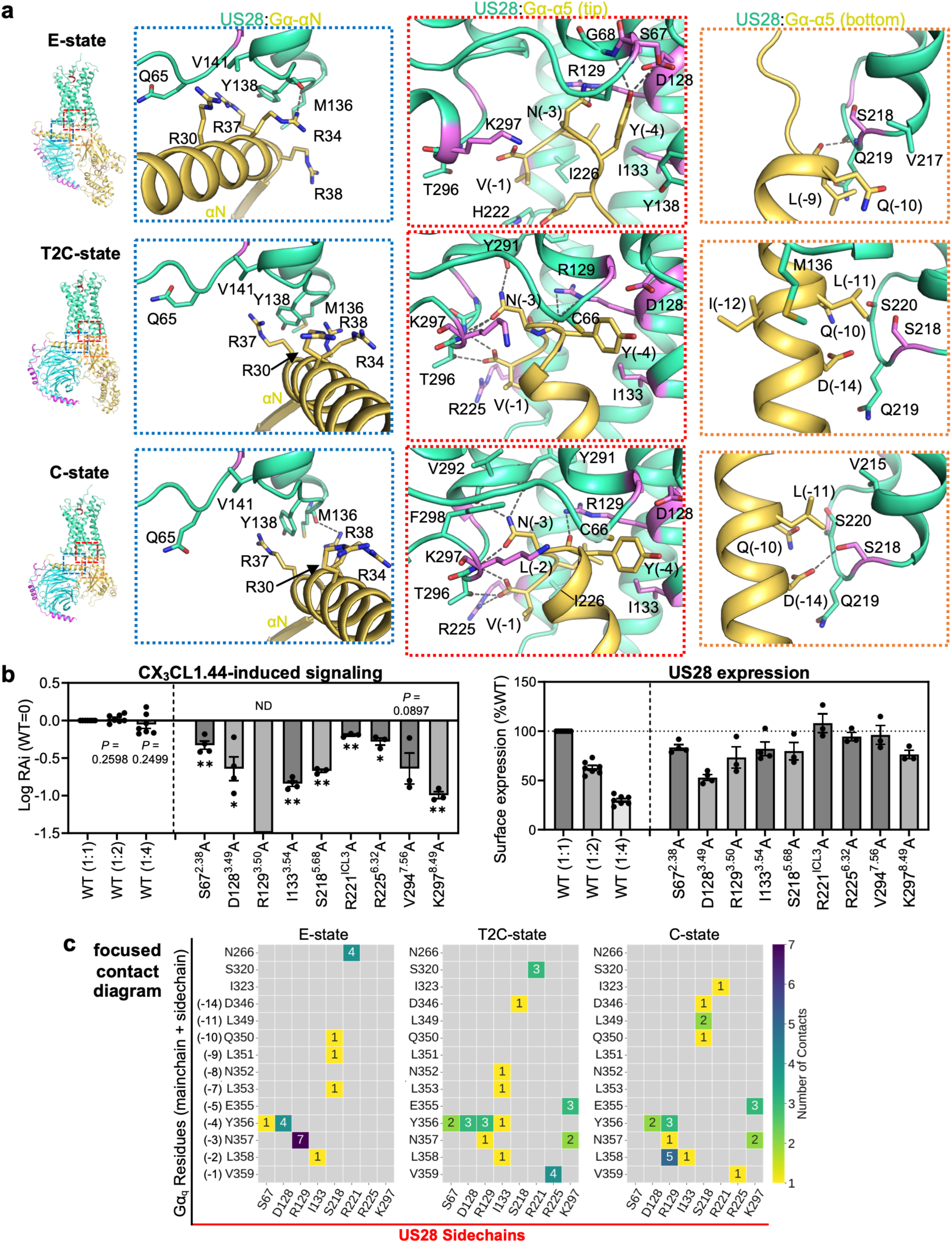
State-dependent interface rearrangements and functional assessments. **a**, Detailed molecular interactions at the US28-G_q_ interface across the E-state (top), T2C-state (middle), and C-state (bottom). Interactions are organized by the involved regions: US28:Gα-αN helix (left), US28:Gα-α5 tip (center), and US28:Gα-α5 bottom (right). Key residues are labeled and shown as sticks. Polar interactions (hydrogen bonds or salt bridges) are indicated by dashed lines. US28 residues selected for mutation are shown in purple. **b**, Functional analysis of the US28-G_q_ interface via site-directed mutagenesis. Left: G_q_ activation by US28Δ300 variants, measured by NanoBiT assay upon CX_3_CL1.44 stimulation. Signaling data are presented as the logarithm of the relative intrinsic activity (LogRAi), normalized to the wild-type (WT) response (WT=0). Right: Cell surface expression of the corresponding US28Δ300 variants, quantified by flow cytometry and normalized to WT (100%). Different dilutions of the WT construct (1:1, 1:2, 1:4) were included as controls for both assays. In both panels, bar colors are shaded according to the relative cell surface expression level. Data represent the mean ± SEM from at least three biologically independent experiments (n = 7 for WT controls; n = 3 or 4 for mutants) each performed with technical triplicate. Analysis of the WT dilution series (1:2 and 1:4) showed no significant difference in LogRAi compared to the WT(1:1) theoretical baseline of 0 (*P* = 0.2598 and *P* = 0.2499, respectively), indicating that the signal is saturated within this expression range. Nevertheless, for a rigorous comparison, variants were stratified into WT(1:1)- or WT(1:2)-matched groups based on their expression levels (see Methods for details). Statistical significance of signaling (LogRAi) was determined by comparing variants to their respective expression-matched WT controls: a two-tailed one-sample *t*-test for the WT(1:1) match or a two-tailed unpaired *t*-test with Welch’s correction for the WT(1:2) match. The R129^3.50^A variant showed completely abolished signaling and was excluded from statistical analysis (ND, not determined). Unadjusted *P*-values from *a priori* planned comparisons were utilized. \**P* < 0.05, \*\**P* < 0.01 versus the respective expression-matched WT control. For the V294^7.56^A variant, the exact P-value (*P* = 0.0897) is displayed to highlight a qualitative trend toward decreased signaling. Comprehensive summary is presented as Supplementary Table 3. **c**, Focused contact diagram of state-dependent interactions at the US28-G_q_ interface. The US28 residues shown correspond to those evaluated in the functional assay (panel b), except for Val294^7.56^, which acts allosterically. Color intensity indicates the number of atomic contacts. Full contact maps are provided in Supplementary Fig. 7.

The engagement of US28 with the Gα_q_-αN helix shifts notably across the states, moving from interactions primarily involving the more N-terminal region of αN in the E-state to the more C-terminal region in the T2C- and C-states (Fig. 4a left and Supplementary Fig. 7). However, the major rearrangement involves the formation and rotation of the α5 helix at the Gα_q_ C-terminus, a key element initiating G protein activation. This movement pivots around the C-terminal Val(−1) residue, which serves as a stationary anchor point stabilized by US28-H8 across all states (Fig. 2c,d and Fig. 4a center) (where −1 is the extreme C-terminus; preceding positions are −2, −3, etc.).

The interactions of the preceding residues are more state-dependent, and the differences propagate as they move away from the depth of the binding pocket. For example, the Asn(−3) and Tyr(−4) sidechains undergo marked reorientations: the former shifts from relatively limited contacts in the E-state to extensive interaction networks involving different receptor elements in the C-state, while the latter transitions from a hydrogen-bond driven interaction with the bottom of TM2 to a van der Waals dominated interaction with TM7 (Fig. 4a center). Similarly, the engagement at the edge of the intracellular pocket undergoes notable rearrangement. In the E-state, the interface is relatively shallow. In the T2C- and C-states, however, the formation of the α-helix draws the C-terminal segment of Gα_q_ deeper into the intracellular cleft near Asp(−14), facilitating numerous new hydrophobic and polar interactions that stabilize the fully engaged complex (Fig. 4a). The complete contact lists are provided in Supplementary Fig. 7, and a more detailed structural analysis of these interactions is available in the Supplementary Discussion.

### Functional assessments of the activation intermediates

To further assess the relevance of each state during G_q_ activation, we probed the effect of single-point US28 mutations at the Gα_q_ interface (Fig. 4b and Supplementary Fig. 8a). To systematically evaluate this multi-step process, we selected mutational targets based on specific structural hypotheses, designing them to probe persistent anchor points among all three states, state-dependent contacts that form or break during activation, and allosteric stabilization networks (Supplementary Table 3). Using the NanoBiT-G_q_ activation assay, which measures Gα_q_-PLCβ association upon G_q_-coupled GPCR ligand stimulation^31^, we assessed CX_3_CL1.44-induced G_q_ signaling for the near full-length US28 (residues 2-354) with an N-terminal FLAG tag. This construct produced a modest ligand-induced G_q_ activation response, likely owing to the constitutive receptor internalization^32^, which limited the dynamic range for quantitative comparison. We therefore used US28Δ300 (ref^33^), a C-terminal truncated construct of US28 (residues 2-300), for the subsequent functional study. This truncation increased surface receptor expression and ligand response, allowing for a more quantitative comparison of G_q_ signaling among receptor mutants.

Our mutagenesis analysis provided complementary data consistent with distinct roles for residues at the G protein interface, supporting a stepwise activation mechanism (Fig. 4a-c). The most substantial effect was observed with the R129^3.50^A mutant, which targets the central arginine of the DR^3.50^Y motif at the core of the G protein interface (superscript refers to Ballesteros-Weinstein numbering). This mutation completely abolished G_q_ activation in response to CX_3_CL1.44. The Arg129^3.50^ sidechain consistently interacts with the Gα_q_ C-terminus across all observed states, from the E-state to the C-state (Fig. 4a center and Fig. 4c). This suggests a dual role of Arg129^3.50^, which not only acts as the primary anchor for the G protein but is also essential for stabilizing the receptor’s active conformation, making it indispensable for signaling^33,34^. Further highlighting the importance of this motif, the D128^3.49^A mutant also caused a substantial loss of G_q_ activation, as its sidechain maintains an interaction with Tyr(−4) of the Gα_q_ C-terminus across all states (Fig. 4a center and Fig. 4c).

Several mutations identified residues that provide continuous structural support throughout activation. The S218^5.68^A mutant, for instance, caused a significant loss of ligand-induced signaling by targeting a residue that maintains persistent interactions across all states (Fig. 4a right and Fig. 4c). By contrast, the V294^7.56^A mutant highlights the importance of allosteric stabilization. Although Val294^7.56^ does not directly interact with Gα_q_, it packs against Val229^6.36^ at the bottom of transmembrane helix (TM) 6 (Supplementary Fig. 8b). Its mutation therefore only resulted in a qualitative trend toward reduced signaling (*P* = 0.0897) by disrupting the receptor’s ability to maintain a fully active conformation.

Other mutants were selected to probe more dynamic behaviors, further linking specific structural states to function. The K297^8.49^A mutant, which had one of the most substantial impacts on signaling, provides a clear example. The interaction involving Lys297^8.49^ is absent in the initial E-state but forms upon reaching the T2C-state and is maintained in the final C-state (Fig. 4a center and Fig. 4c). Therefore, the severe impairment of signaling by the K297^8.49^A mutation is consistent with the destabilization of the T2C- and C-states, possibly by inhibiting the E-to-T2C transition.

Conversely, the R221^ICL3^A mutation slightly impaired signaling by targeting a residue whose contribution is greatest during the initial E-state and diminishes as activation proceeds (Fig. 4c and Supplementary Fig. 8c). Likewise, the slightly reduced signaling observed with the S67^2.38^A mutant likely underscores the importance of the early activation steps, as this residue forms more extensive interactions with G_q_ in the E- and T2C-states than the C-state (Fig. 4a center and Fig. 4c). The functional relevance of the T2C intermediate was further assessed by the I133^3.54^A and R225^6.32^A mutants (Fig. 4a center and Fig. 4c). These were designed to disrupt a transiently increased interface more selectively in the T2C-state, and both resulted in signal reduction.

Collectively, while these cellular assays cannot isolate individual intermediate steps, the structure-guided mutagenesis data are consistent with the hypothesis that these three states represent the physiological G_q_ activation pathway.

### A stepwise allosteric cascade drives GDP release

In contrast to the constitutively active conformation of US28, our structures support a stepwise G_q_ activation mechanism that progresses from the inactive, resting state, through the key intermediate E- and T2C-states, to the final nucleotide-free C-state. Each state provides a distinct snapshot of the conformational changes in G_q_ required to unlock its nucleotide-binding pocket and release GDP (Fig. 5, 6, and Supplementary Fig. 9). Based on structural comparison, we propose the following mechanism.

**Fig. 5.**
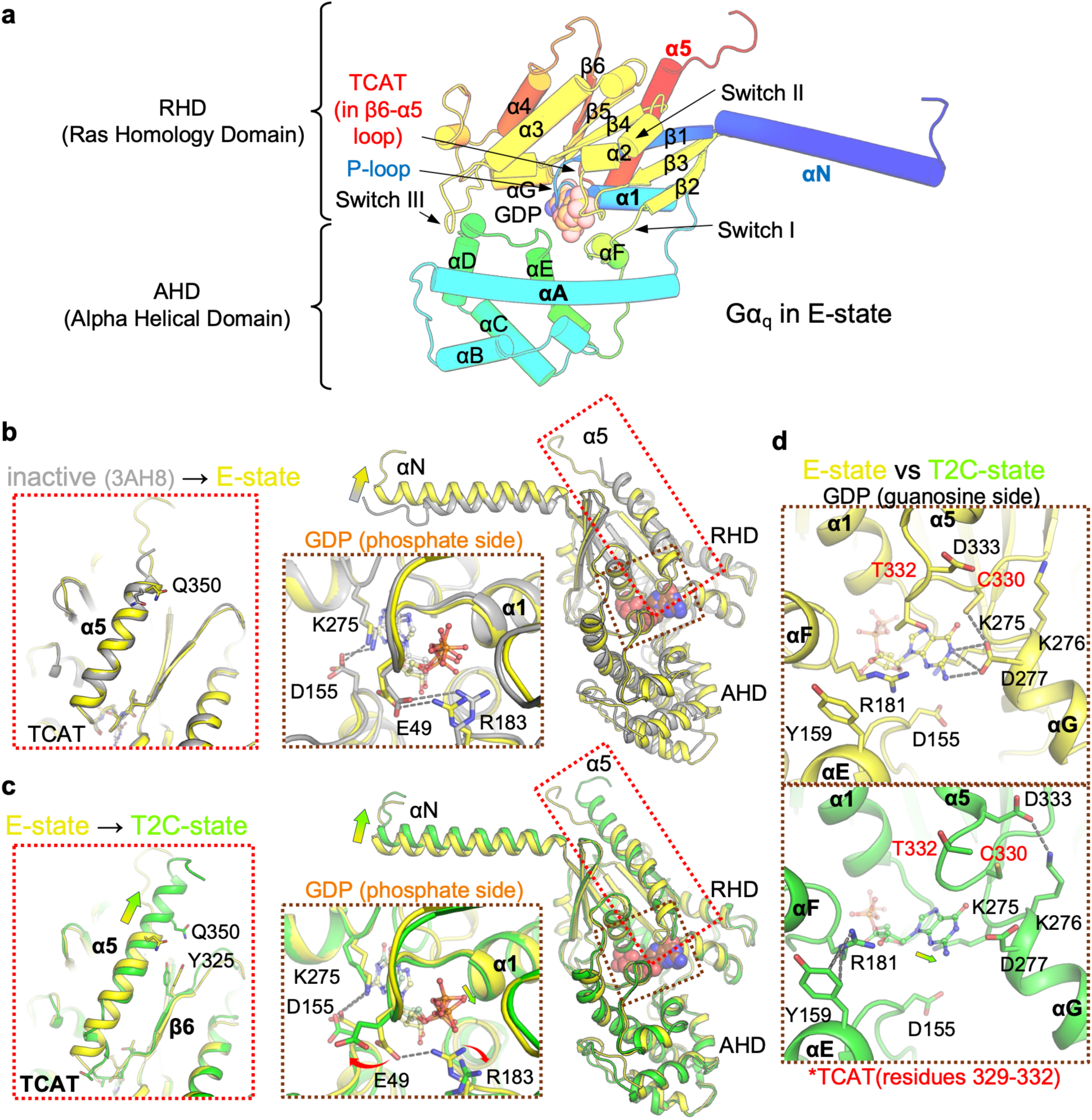
Allosteric cascade driving GDP release: from the inactive to T2C-state via E-state. Structural comparison of Gα_q_ showing the initial conformational changes leading to Gα_q_ pre-activation. Structures are aligned based on the RHD. **a**, Structural elements of Gα_q_ protein. Gα_q_ from the E-state is shown as cartoon figure and labeled with key secondary or functional structure elements. The protein is colored by a spectral gradient from the N-terminus (blue) to the C-terminus (red). **b**, Transition from the inactive, GDP-bound Gα_q_ (PDB: 3AH8, gray) to the E-state (yellow). Insets highlight the TCAT motif and α5 (left inset), and the nucleotide-binding pocket with the Glu49-Arg183 salt bridge (middle inset). GDP is shown as ball-and-stick. **c**, Transition from the E-state (yellow) to the T2C-state (green). Deeper insertion of the α5 helix perturbs the TCAT motif (left inset). Crucially, the Arg183 sidechain (Switch I) flips outward, breaking the Glu49 (P-loop)-Arg183 (Switch I) salt bridge (middle inset). **d**, Comparison of the guanosine binding pocket in Gα_q_ between the E- (yellow) and T2C-states (green). Remodeling of the TCAT motif (including Cys330 and Thr332) and repositioning of the GDP molecule disrupt key interactions shielding the guanosine moiety, thereby lowering the affinity for GDP and destabilizing the RHD-AHD packing.

**Fig. 6.**
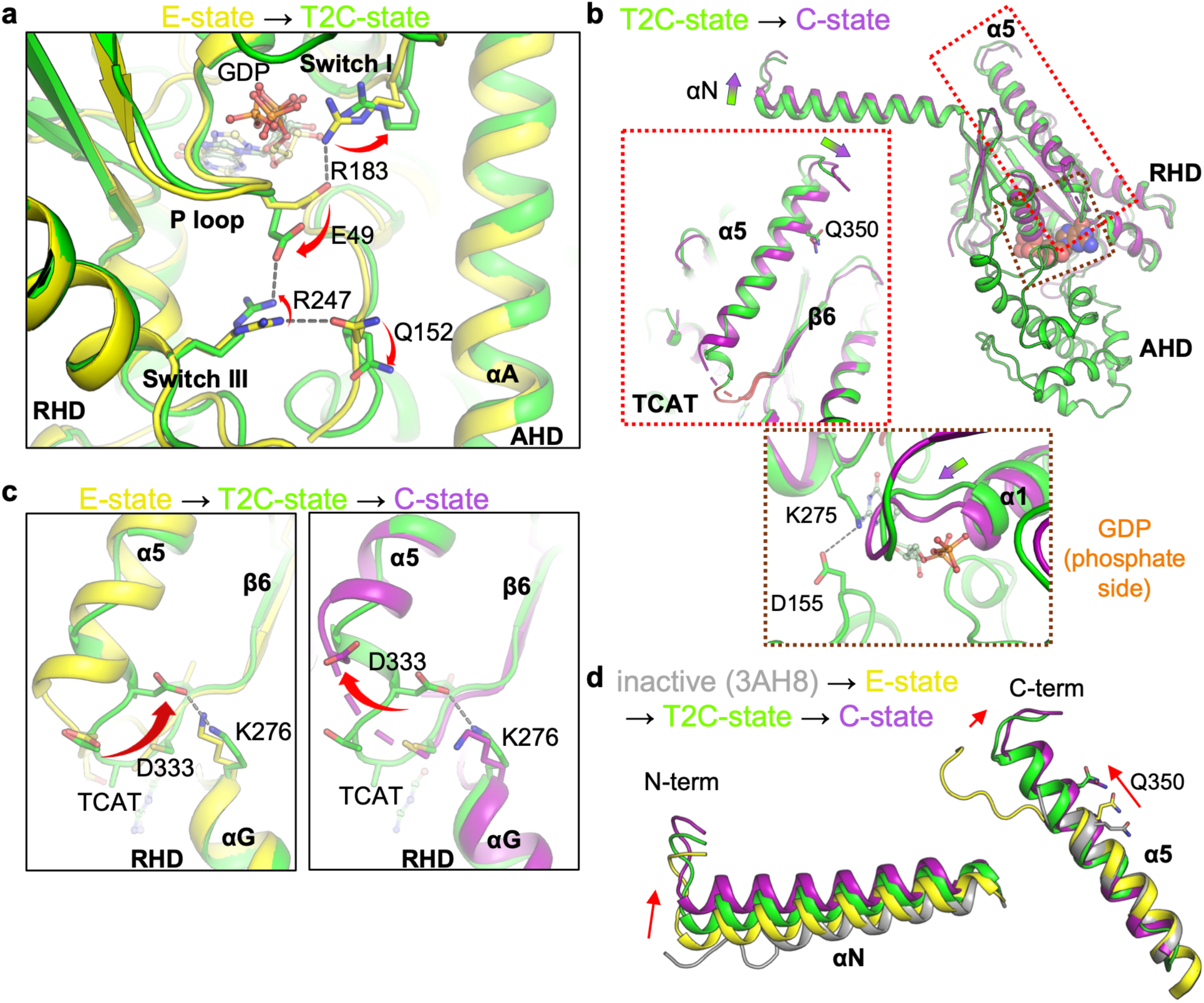
Allosteric cascade driving GDP release: from the E-state to C-state via T2C-state. Structural comparison of Gα_q_ showing the stepwise conformational changes leading to nucleotide release. Structures are aligned based on the RHD. **a**, Structural details of the allosteric relay in Gα_q_ from the E-state to the T2C-state, focusing on the salt bridge relay. Glu49 forms a new interaction with Arg247 (Switch III), which consequently breaks the interaction between Arg247 and Gln152 (AHD). **b**, Transition from the T2C-state (green) to the C-state (magenta). Completion and linearization of the α5 helix (left inset) drives the displacement of the α1 helix (bottom, arrow). This displacement physically expels GDP and promotes the separation of the AHD from the RHD (bottom inset). **c**, Close-up view of the TCAT loop during transition from E-state (yellow) to T2C-state (green) to C-state (purple). Movement of Asp333 alters the conformation of the TCAT in each state. The TCAT motif eventually moves away from the guanosine moiety and becomes disordered, reducing its affinity for GDP. **d**, Relative conformational changes of α5 and αN amongst the inactive, E-, T2C-, and C-states Gα_q_, showing the consistent clockwise movement when viewed down from the extracellular side.

In the E-state, the G protein is poised for activation, with the C-terminal region of Gα_q_ partially structured to engage the intracellular core of activated US28 (Fig. 5b). Although the G protein’s overall conformation resembles its inactive, GDP-bound state (PDB: 3AH8)^35^, there are crucial distinctions. The packing between the AHD and RHD is slightly loosened by a hinge-like swing of the AHD toward the RHD’s α5 helix (Supplementary Fig. 9a). When the structures are aligned on the RHD, this rotational displacement of the AHD is characterized by a 2.5 Å shift at the C-terminus of α-helix A (αA), resulting in an overall Cα RMSD of 2.2 Å between the AHDs (113 atoms). The conformation of the AHD itself is essentially maintained, with an internal Cα RMSD of 0.54 Å. This conformational change, also seen in G_11_ (Supplementary Fig. 9b), is associated with an altered interdomain lock in G_q_. While Switch III engages both αA and the αD-αE loop of the AHD in the inactive state, its interaction in the E-state is maintained only through the αD-αE loop. The E-state conformation is reminiscent of intermediate states observed during G protein dissociation following GTP binding^11^.

The transition from the E- to T2C-state triggers a sequence of allosteric rearrangements, initiated as the Gα_q_ α5 helix is almost fully formed (Fig. 5c). The upward movement of the α5 helix directly perturbs the TCAT-motif containing β6-α5 loop and slightly draws the β6 strand upward primarily via an interaction with Tyr325. This perturbation flips the Cys330 and Thr332 sidechains away from the GDP cage and repositions the GDP molecule by pulling its guanosine moiety toward the αG side (Fig. 5d). This action results in a primary displacement of the guanosine base, while the phosphate group shifts slightly downward to the AHD side. This overall displacement repels the Asp277 and Arg181 sidechains (Fig. 5d bottom), increasing the solvent accessibility of the guanosine moiety, and also pushes the Arg183 sidechain at the phosphate side (Fig. 5c center). These downstream movements subsequently break the salt bridge between Glu49 in the P-loop and Arg183 in the Switch I region that had previously shielded the GDP molecule. While the disruption of the Glu49-Arg183 salt bridge has been suggested to be the rate-limiting step in G_q_ and G_s_ activation^15,16,36^, we view it as a consequence of α5 engagement. The Glu49 flip further exposes the phosphate group of GDP to solvent and enables formation of a new salt bridge between Glu49 and Arg247 in the Switch III loop (Fig. 6a). The formation of the new Glu49-Arg247 bridge concurrently breaks a critical interaction between Arg247 (Switch III) and Gln152 (AHD) (Fig. 6a). This sequential breakage and formation of ionic interactions constitute a “salt bridge relay” that transmits the allosteric signal from Switch I to the AHD interface. As a result, the AHD-RHD interface is destabilized, and the Switch III loop gains flexibility, collectively unlocking the G protein for GDP release.

Finally, in the C-state, these preparatory changes trigger the expulsion of GDP (Fig. 6b). As the bending tension within the α5 helix is released, its receptor-distal region simultaneously becomes fully ordered (Fig. 6c,d). This process completes the α5 helix in a straightened conformation that, in turn, perturbs the TCAT motif and triggers the displacement of the α1 helix (Fig. 6b, c). The displaced α1 helix creates a steric clash that physically expels GDP and subsequently pushes the AHD away, separating it from the RHD and minimizing GDP affinity (Fig. 6b). Untethered from the RHD, the AHD becomes highly dynamic, stochastically sampling a wide ensemble of conformations and thereby generally precluding definitive structural modeling of its position. However, we also captured a distinct nucleotide-free state, C’-state, in which a transiently closed AHD conformation appears slightly stabilized (Supplementary Fig. 5c).

During this activation process, the TCAT motif is gradually perturbed (Fig. 6c). Hydrogen bond formation between Asp333 and Lys276 in the T2C-state appears to ‘springload’ the α5 helix, which becomes distorted at the base. Upon transition to the C-state, this hydrogen bond is released, resulting in re-formation of the α5 helix base and disordering of the β6-α5 loop. Concurrently, the αN helix progressively draws closer to the α5 helix (Fig. 6d), which perturbs the downstream β2-strand, P-loop, and α1 helix. Notably, our structures confirm the inhibitory mechanism of the G_q/11_-selective compound YM-254890^35,37,38^. This compound wedges itself between the α1 helix and the β2-strand, thereby preventing the critical displacement of the α1 helix required to physically expel GDP. This steric hindrance mechanism complements its function as a “molecular glue” that stabilizes the interface between the RHD and AHD^38^. Therefore, its potent inhibition arises from this dual mechanism of physically impeding essential conformational rearrangements and gluing the G protein in an inactive state.

## Discussion

The activation of heterotrimeric G proteins by GPCRs involves a complex series of conformational changes culminating in GDP release. While the nucleotide-free endpoint has been extensively characterized, the dynamic trajectory leading to it has remained elusive, with insights often relying on computational approaches^14,15^. By leveraging the properties of the viral GPCR US28, we have captured a series of high-resolution snapshots detailing the stepwise activation of G_q_ (Fig. 1). Furthermore, this study revealed the structural basis of G protein subtype selectivity and biased chemokine agonism (Supplementary Discussion and Supplementary Fig. 10, 11).

A pivotal finding of this study is the visualization of the T2C-state, a plausible on-pathway intermediate where the Gα_q_ C-terminus is fully engaged with the receptor while GDP remains bound. This structure provides the direct structural evidence for a key allosteric event: the disruption of the critical Glu49 (P-loop)-Arg183 (Switch I) salt bridge. Previous MD simulations predicted this event to be the rate-limiting step in spontaneous G_q_ activation^15^ but lacked experimental validation. Our T2C-state structure reveals that the insertion of the α5 helix coincides with this critical event before the large-scale AHD opening.

This mechanism aligns closely with the activation process proposed for G_s_. Extensive studies of the β2AR-G_s_ complex identified the disruption of the analogous salt bridge (Glu50- Arg201 in G_s_) as the critical event for GDP release^16,25^. The R201C mutation, which abolishes this interaction, renders G_s_ constitutively active^36^. Furthermore, this Glu-Arg residue pair is broadly conserved among the Gα_s_, Gα_i_, Gα_q_, and Gα_12/13_ subfamilies, with Gα_z_ (Asn-Arg) being the sole exception among the 16 human Gα subunits^39^. Thus, our direct structural observation of this conserved intermediate step in G_q_ provides strong evidence for a unified activation principle across different G protein families, in which the receptor-induced disruption of this specific salt bridge serves as the universal step toward unlocking the G protein.

The T2C-state reveals precisely how this conserved allosteric relay progresses. Rearrangement of the RHD, driven by α5 insertion, initiates a critical salt bridge relay: Glu49 flips outward to form a new interaction with Arg247 (Switch III), which in turn breaks the interaction between Arg247 and Gln152 (AHD) (Fig. 6a). This destabilizes the AHD-RHD interface and increases GDP solvent accessibility. Crucially, our structures, combined with MD simulations, demonstrate that the unlocking of the nucleotide-binding pocket is a distinct event that precedes the large-scale opening of the AHD. This mechanism decouples the initial allosteric perturbation at the nucleotide pocket from the final GDP expulsion step, which is subsequently driven by the displacement of the α1 helix in the C-state.

The comprehensive view of the activation trajectory also allows us to contextualize these findings within the full G protein cycle. The E-to-C transition involves an approximately 33° rotation of G_q_ relative to US28. Interestingly, this rotational movement during activation is similar to the trajectory observed during the GTP-induced dissociation of the β2AR-G_s_ complex^11^. However, this differs from the “sliding” dissociation trajectory observed for the μ-opioid receptor (μOR)-G_i_ complex^17^.

These differing dissociation trajectories, rotation for G_s_/G_q_ versus sliding for G_i_, do not contradict the unified activation mechanism. Rather, they suggest a multi-layered process where a conserved core mechanism (salt-bridge disruption) during activation is complemented by family-specific dissociation dynamics. We speculate that the specific mode of dissociation might fine-tune the kinetics of signaling. For example, the rotational movement observed in G_q_/G_s_ might facilitate rapid GTP binding and dissociation, enabling fast signal amplification, whereas the sliding mechanism in G_i_ might favor different kinetic properties tailored to the specific receptor-G protein interface.

While finalizing this manuscript, a landmark study unveiling the agonist-induced GDP dissociation in the μOR-G_i_ system was published by Khan *et al.*^40^. The structures are complementary to, yet mechanistically distinct from our intermediate states (Fig. 7). Their “Latent state” is similar to our E-state, but the α5 helix is positioned at the outside of the transducer binding pocket and sticks at the lateral surface of the intracellular loops (Fig. 7a), as observed for the GTP-bound μOR-G_i_ complex “G-ACT-3“^17^. Likewise, their “Engaged state” corresponds to our T2C-state but displays a fully formed, nucleotide-free-like α5 helix (Fig. 7b). Despite these conformational differences, the critical disruption of the P-loop/Switch I salt bridge is conserved in both intermediate states (Fig. 7c, d), although Arg178 in the Engaged μOR-G_i_ complex inserts into the magnesium-binding site (Fig. 7d, rightmost panel). Furthermore, our GDP-bound/AHD-open state in the MD simulations (Fig. 3b) is reminiscent of their “Primed” state, which immediately precedes GDP release.

**Fig. 7.**
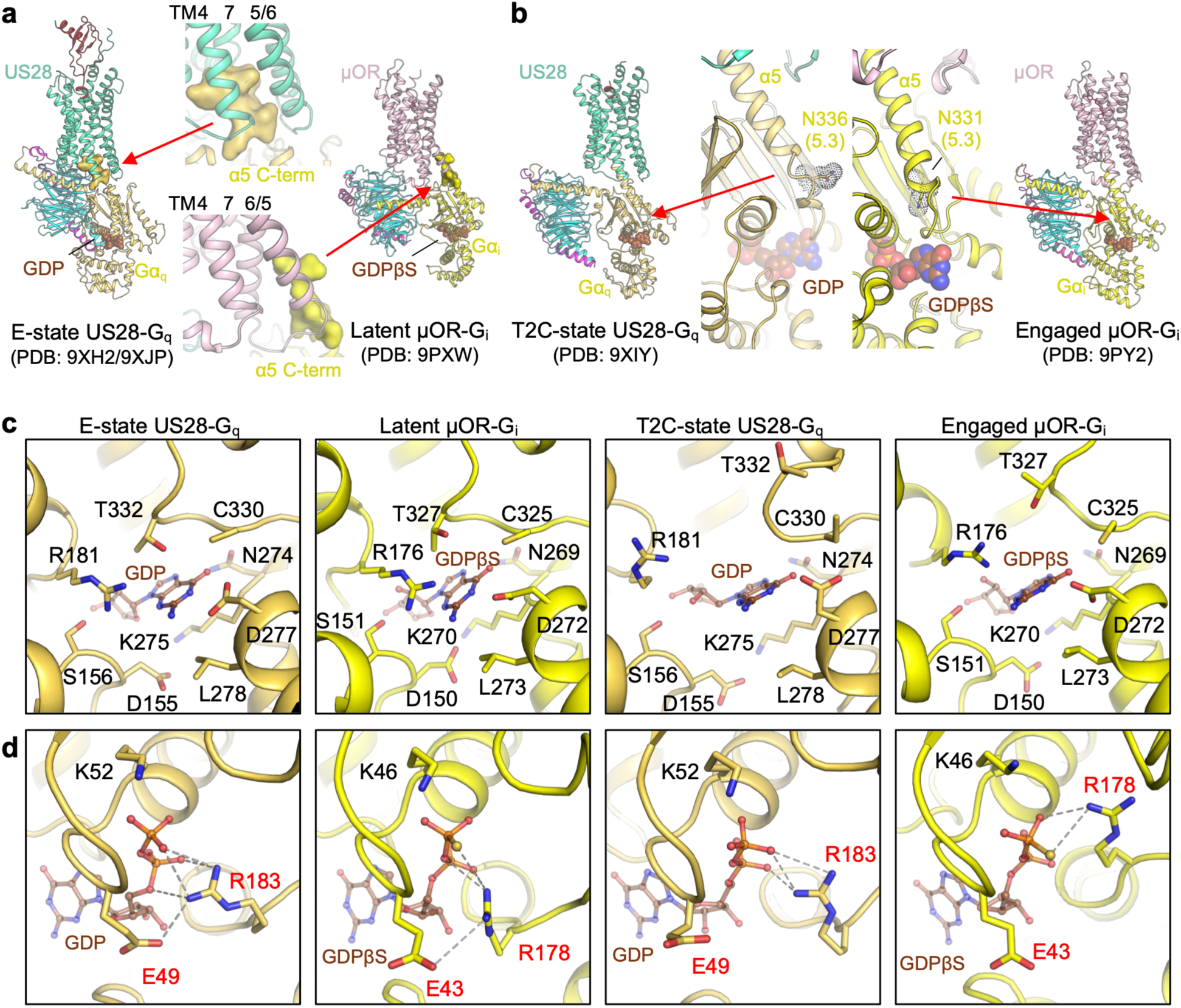
Structural comparison of plausible G protein activation intermediates between the US28-G_q_ and μOR-G_i_ complexes. **a**, Comparison of the initial encounter states: the US28-G_q_ E-state (left) and the naloxone-bound Latent μOR-G_i_ (right; PDB: 9PXW). In the US28-G_q_ E-state, the C-terminal region of the Gα_q_-α5 is inserted into the intracellular cavity of the receptor. In contrast, the α5 helix in the Latent μOR-G_i_ is positioned outside the transducer-binding pocket, packing against the lateral surface of the intracellular loops. The C-terminal seven residues of the α5 helix (Leu(−7) to Val(−1) for Gα_q_, and Leu(−7) to Phe(−1) for Gα_i1_) are shown in a surface representation. **b**, Comparison of the subsequent intermediate states prior to GDP release: the US28-G_q_ T2C-state (left) and the Engaged loperamide-bound μOR-G_i_ (right; PDB: 9PY2). Both states capture a transient phase where the α5 helix is deeply inserted into the receptor core while GDP/GDPβS remains bound. However, the membrane-distal segment of the α5 helix remains broken in the US28-G_q_ T2C-state. Conversely, the μOR-G_i_ Engaged state displays a fully formed, nucleotide-free-like α5 helix. In panels a and b, the structures are superimposed on the receptors. High-resolution μOR-G_i_ structures bound to different ligands (naloxone and loperamide) were utilized to ensure the most accurate structural comparison. **c, d,** Close-up views of the nucleotide-binding pocket. c, Comparison of the guanosine-binding site across the intermediate states. Key pocket residues of the E-state and the corresponding residues in the other states are shown as sticks. **d,** Conserved intermediate event at the phosphate-binding side. Despite differences in the initial engagement modes and α5 conformations, both intermediate trajectories feature the critical disruption of the conserved salt bridge between the P-loop glutamate (Glu49 in Gα_q_; Glu43 in Gα_i1_) and the Switch I arginine (Arg183 in Gα_q_; Arg178 in Gα_i1_).

The apparent distinctions may reflect both the intrinsic differences between the receptor-G protein pairs (μOR-G_i_ vs. US28-G_q_) and the experimental methodologies employed; their states were captured using a “re-bound GDP” protocol, which likely represents a slightly different equilibrium state, while our structures were captured using a “GDP removal” protocol without nucleotide re-addition. Nevertheless, it is noteworthy that both studies identify the disruption of the conserved Glu-Arg salt bridge as a key structural change for unlocking the G protein. This convergence, despite the distinct receptor systems and intermediate states observed, strongly reinforces our conclusion that the receptor-induced breakage of this specific ionic lock is a key universal event initiating G protein activation across different families.

### Conclusion and limitations of the study

While our integrated approach provides a compelling model for stepwise G protein activation, several caveats temper our conclusions. The structures were determined in detergent micelles with a stabilizing scFv16 fragment, which may shift the equilibrium of transient states; however, MD simulations in a lipid bilayer without scFv16 maintained a stable T2C-state capable of bidirectional transitions mainly toward the C-state (with 6 out of 16 simulations transitioning toward the C-state and 1 out of 16 toward the E-state), indicating that it is a physically viable conformation rather than a micelle-induced artifact. Admittedly, exploring the simulated trajectory solely from the T2C-state inherently limits the identification of other potential intermediates or pathways. Nevertheless, the experimental capture of the C’-state may offer a valuable structural reference for such alternative conformations. The chimeric Gα_q_ (Gα_qiN18_) used for scFv16 binding introduces a non-native element, although this substitution is strictly confined to the first 18 N-terminal residues, which are spatially distant from the receptor interface. Furthermore, while our cellular mutagenesis data are consistent with the structural states, endpoint assays cannot resolve individual kinetic steps. Defining the precise temporal sequence would require real-time single-molecule or time-resolved assays, which remain technically challenging here. The proposed mechanism is therefore best viewed as a plausible, structurally grounded model. Finally, we cannot exclude the existence of alternative pathways, which might be favored in distinct environments.

Within these bounds, our study provides a high-resolution structural view of the G_q_ activation trajectory catalyzed by a GPCR. The stepwise mechanism, in particular the conserved allosteric relay preceding AHD opening, supports a unified model for GPCR-mediated G protein activation.

## Methods

### Preparation of CX_3_CL1.44-US28-G_qiN18_-scFv16 complex

CX_3_CL1.44, an engineered version of the chemokine domain of CX_3_CL1, and full-length US28 were expressed separately with the BacMam system using HEK293S GnTI^-^ cells, and the CX_3_CL1.44-US28 complex was purified as described^13^. Briefly, C-terminal Fc-tagged CX_3_CL1.44 was first captured from culture medium by Protein A Sepharose (GE Healthcare/Cytiva). Cell membranes containing N-terminally FLAG-tagged US28 were isolated and solubilized in HEPES-Buffered Saline (HBS, 10 mM HEPES-Na pH 7.2 and 150 mM NaCl) containing 1% (w/v) LMNG/0.2% (w/v) CHS, and FLAG-US28 was captured by CX_3_CL1.44-Fc immobilized on Protein A Sepharose. The resin was washed with HBS containing 0.1% (w/v) LMNG/0.02% (w/v) CHS, and Fc-tag was cleaved off by 3C protease on Protein A column. The eluate containing CX_3_CL1.44-US28 was purified by the anti-FLAG M1 Sepharose with decreased detergent concentration to 0.01% (w/v) LMNG/0.002% (w/v) CHS. The eluted complex was further purified over a size-exclusion chromatography (SEC) column Superose 6 10/300 GL (GE Healthcare/Cytiva) equilibrated with HBS containing 0.005% (w/v) LMNG/0.0005% (w/v) CHS.

To prepare G_qiN18_, Gα_qiN18_ was designed similarly to Gα_11iN18_ (ref^27^) by replacing the N-terminal 18 amino acid residues with the corresponding region of Gα_i1_ for scFv16 binding to assist in minimizing dissociation after CX_3_CL1.44-US28-G_q_ complex formation and to aid in cryo-EM image processing. Gα_qiN18_ was co-expressed with 6xHis-tagged Gβ_1_ and Gγ_2_ in *Trichoplusia ni* cells (Expression Systems), and purified using Ni-NTA agarose (Qiagen) as a GDP-bound heterotrimer in the presence of 10 μM GDP as previously described^41^. The scFv16 used for the structural study was secreted from *Trichoplusia ni* cells and purified using Ni-NTA agarose^13,27^.

To form the CX_3_CL1.44-US28-G_qiN18_ complex, 1.2 molar equivalents of G_qiN18_ were used for the coupling reaction with CX_3_CL1.44-US28 in the presence of 1% (w/v) LMNG/0.1% (w/v) GDN/0.1% (w/v) CHS. Residual GDP in reaction buffer was hydrolyzed with apyrase (NEB), reducing the chance of GDP rebinding from the buffer solution. The complex was then purified over a FLAG column to remove excess G_qiN18_ and eluted in HBS containing 0.001% (w/v) LMNG/0.001% (w/v) GDN/0.0001% (w/v) CHS, 5 mM EDTA and 0.2 mg/ml FLAG peptide. 0.1 mM TCEP was added immediately after protein elution. 1.2 molar equivalents of scFv16 were added to the FLAG eluate containing CX_3_CL1.44-US28-G_qiN18_, and the mixture was incubated on ice for 2 hrs. The mixtures were loaded onto a Superdex 200 10/300 GL equilibrated with HBS containing 0.001% (w/v) LMNG, 0.001% (w/v) GDN, 0.0001% (w/v) CHS, and 0.1 mM TCEP (SEC buffer), and the peak fractions were concentrated to ∼20 mg/ml.

### Cryo-EM grid preparation and data collection

To prepare the cryo-EM specimen, 10% volume of the SEC buffer containing either 0.1% (w/v) (1H, 1H, 2H, 2H-Perfluorooctyl)-β-D-Maltopyranoside (fOM, Anatrace) or 10 mM 3-[(3-cholamidopropyl)dimethylammonio]-2-hydroxy-1-propanesulfonate (CHAPSO, Sigma) was added to the sample. 3 µL of each mixture was applied to glow-discharged 300 mesh gold grids (Quantifoil R1.2/1.3). Excess sample was blotted to a filter paper for 3 sec before plunge-freezing using a Leica EM GP (Leica Microsystems) at 16°C and 95% humidity. The cryo-EM movies were collected on a Titan Krios operated at 300 kV and equipped with a Gatan K3 camera and BioQuantum imaging filter in super-resolution mode. Magnification was set to 105,000x, corresponding to a non-super resolution pixel size of 0.8677 Å. Movies were recorded for a total of 2.5 sec with 0.05 sec exposure per frame at an exposure rate of ∼20 electrons/pixel/sec and the nominal defocus range between −0.8 and −2.0 μm, using SerialEM^42^ with beam-image shift to collect 9 images from 9 holes per stage shift and focus.

### Cryo-EM data processing

For the raw movies, beam-induced motion correction and estimation of patch contrast transfer function (CTF) parameters were performed using cryoSPARC version 2 Live^43^. Low-quality micrographs were discarded and 3,255 and 8,518 movies from the cryo-EM samples prepared with fOM and CHAPSO, respectively, were used for the following data processing in the main interface of cryoSPARC version 4 (ref^43^), which is summarized in Supplementary Fig. 1d and 2.

After patch motion correction and CTF estimation, particles were picked iteratively with multiple methods (Gaussian picking, template-based picking, and Topaz picking^44^) combined with 2D classifications, which yielded 8,400,483 initial protein particles extracted with 300/64 pixel binning (4.067 Å/pixel). Further 2D classifications were performed to discard low-quality particles and to select classes with one or two US28 molecules each with a G protein bound. The particles were re-extracted with 352/160 binning (1.909 Å/pixel) for additional 2D classification, multi-class ab-initio 3D reconstruction, non-uniform refinement (NU-refine), and 3D classification.

The class containing a single US28 molecule was first separated into E-state-like (812,733 particles) and C-state-like (1,131,098 particles) populations, which were subjected to re-extraction with 352/300 binning (1.018 Å/pixel). For the E-state like particles, additional 2D and 3D classifications identified the highest-quality particles set, which was subjected to NU-refine, CTF refinement, reference-based motion correction (RBMC), further 3D and 2D classifications, and local NU-refine. The selected 193,867 particles yielded the E-state map with 2.6 Å nominal-resolution determined by gold-standard FSC with the 0.143 criterion, albeit with poor density at the chemokine body (map 1). To improve chemokine density, another round of 3-class 3D classification was performed with the mask covering CX_3_CL1-US28 followed by local NU-refine. The E-state class with the best chemokine density was refined with 64,658 particles to 2.7 Å nominal-resolution (map 2).

The E-state-like class included 186,061 C-state particles, which have closer packing between US28 and G_q_, and exhibit an α-helical structure at the C-terminus of G_q_ typical of GPCR-G protein complexes. This particle set was merged with the C-state like 1,131,098 particles for 3D and 2D classifications, NU-refine, CTF refinement, and RBMC, yielding four C-state classes with AHD separated from RHD (open) and one C-state class with the AHD packed with RHD (closed), albeit with relatively less-defined cryo-EM density. The former C-state classes were essentially identical in the GPCR-G_q_ docking mode with slightly different GPCR-G_q_ angles, which we considered stochastic flexibility. We thus focused on the class that yielded the highest quality map, which generated a 2.6 Å nominal-resolution map with 268,599 particles (map 3). The chemokine core density was improved in the same manner as the E-state to yield a 2.8 Å nominal-resolution map with 89,862 particles (map 4).

The assumed C-state class with partially closed AHD was classified into four 3D classes with the mask covering G_q_-AHD followed by NU-refine, yielding two classes with stably closed AHD and two classes with flexible AHD. One of the AHD-closed classes retains a GDP molecule and was refined to 2.9 Å nominal-resolution with 70,394 particles (map 5-1). The other class lacks clear density around the GDP-binding pocket and was refined to 2.9 Å nominal-resolution with 69,523 particles (map 6-1). As the former appeared the intermediate between the E- and C-states, we termed this state the T2C (Transition-to-Canonical) state, and the other as the C’-state. To maximize interpretability of the maps, we locally refined the US28 region and the AHD/RHD region of Gα_q_ in conjunction with CTF refinement, additional RBMC, and local NU-refine (maps 5-2, 5-3, 6-2, and 6-3). Each map had enhanced local features and was used to guide model building of the complex in the globally refined map.

Map refinement statistics are summarized in Supplementary Table 1. The maps were visualized using UCSF ChimeraX^45^, and local resolution was estimated using Phenix^46^.

### Model building and refinement

A model of Gα_qiN18_ was generated using AlphaFold3^47^ and docked, along with Gβ_1_-Gγ_2_-scFv16 from PDB entry 7RKF, into the E-state map 1 with Phenix Dock in Map. US28 from PDB entry 4XT1 was placed using Phenix EMPlacement^48,49^. The CX_3_CL1.44 N-terminal peptide was interactively built using Coot^50^. The chemokine body of CX_3_CL1 was docked onto this model in E-state map 2 using Coot. The remaining 1:1 complexes were initially docked by the same methods using models of the Gα_qiN18_-Gβ_1_-Gγ_2_-scFv16 (with the AHD removed as appropriate) and US28-CX_3_CL1.44 (full-length or N-terminal peptide) from the E-state for C-state with the N-terminal peptide (map 3), C-state with full-length chemokine (map 4), T2C-state (map 5-1), and C’-state (map 6-1).

Further interactive model building for all structures was performed in Coot using the uniformly sharpened maps and DeepEMhancer-sharpened maps. Real space refinement and ADP refinement were performed in Phenix using secondary structure restraints defined using ksdssp^51^. Model quality was assessed using Molprobity^52^. Model statistics are reported in Supplementary Table 1. Software for model building and refinement was installed using SBGrid^53^. Structural figures were prepared using UCSF ChimeraX and PyMOL (Schrödinger).

### Plasmid construction for functional analysis

Near full-length US28 (residues 2-354) was cloned into the pCAGGS vector^54^ with an N-terminal HA secretion signal peptide, a FLAG tag, and a 16-amino acid flexible linker (MKTIIALSYIFCLVFA-DYKDDDDK-GGSGGGGSGGSSSGGG). For the mutagenesis study, we used a C-terminally truncated US28 construct (residues 2-300, US28Δ300), which was shown to improve cell-surface expression and signal-to-noise ratio in the signaling assay^33^. Structure-guided single point mutations at the interface between US28 and G_q_ were introduced into the pCAGGS US28Δ300 vector using the PrimeSTAR Mutagenesis Basal Kit (Takara).

### NanoBiT G_q_ activation assay

G_q_ activation assays were performed as described previously^31^. Briefly, LgBiT-tagged Gα_q_ and SmBiT-tagged PLCβ2 were co-transfected using polyethylenimine (PEI) Max solution (1 mg/mL; Polysciences) into HEK293A cells (Thermo Fisher Scientific) with a US28 construct along with Gβ_1_, Gγ_2_, and Ric8A. After one-day incubation, the cells were harvested, resuspended in Hank’s Balanced Salt Solution (HBSS) containing 0.01% (w/v) bovine serum albumin (BSA; fatty acid-free grade; SERVA) and 5 mM HEPES (pH 7.4) (assay buffer), reseeded in a white 96-well plate and incubated with 10 µM coelenterazine (Angene) for 2 h. After reading basal luminescence counts, the cells were stimulated with titrated concentrations of CX_3_CL1.44 (Supplementary Fig. 8a), and measured for second luminescence counts. The fold-change values were further normalized to those of vehicle-treated samples and used to plot the G protein dissociation response. G protein activation signals were fitted to a four-parameter sigmoidal concentration-response curve with a constrain of the *HillSlope* parameter to absolute values less than 1.5 using the Prism 10 software (GraphPad Prism). For each replicate experiment, the *Span* parameter (= *Top* – *Bottom*), EC_50_ and *Span*/EC_50_ of the individual US28 mutants were normalized to those of the WT US28Δ300 performed in parallel and logarithmic values of the resulting relative intrinsic activity (LogRAi) were used to calculate ligand response activity of the mutants.

### Flow cytometry

Plasmid transfection was performed according to the same procedure as described in the “NanoBiT G_q_ activation assay” section. The transfected cells were collected and transferred to a 96-well plate and fluorescently labeled with an anti-FLAG epitope (DYKDDDDK) tag monoclonal antibody (Clone 1E6, FujiFilm Wako Pure Chemicals; 10 μg /mL) and a goat anti-mouse IgG secondary antibody conjugated with Alexa Fluor 488 (Thermo Fisher Scientific; 10 μg/mL). Fluorescent intensity was quantified by an CytoFLEX flow cytometer (Beckman Coulter) equipped with a 488-nm laser. The flow cytometry data were analyzed with the FlowJo software (FlowJo). Values of mean fluorescence intensity from approximately 20,000 cells per sample were used for analysis.

### Statistical analysis for functional analysis

To evaluate whether receptor surface expression levels limit the signaling readout, the LogRAi values of the WT dilution series (1:2 and 1:4) were first compared to the WT(1:1) control (theoretical baseline of 0) using a two-tailed one-sample *t*-test. No significant differences were observed (*P* = 0.2598 for 1:2 and *P* = 0.2499 for 1:4), indicating that the signaling response is saturated relative to the WT expression level in this assay system. Thus, within this receptor concentration range, the surface expression level does not affect the LogRAi.

Nevertheless, to align the functional comparison conditions as closely as possible and adopt a rigorous analytical framework, US28 variants were stratified into expression-matched groups based on their closest mean surface expression. Specifically, an absolute expression cut-off of 81.4%, the exact midpoint between the WT(1:1) and WT(1:2) means, was applied as a practical threshold; variants with expression ≥ 81.4% were assigned to the WT(1:1)-matched group, and those with < 81.4% to the WT(1:2)-matched group. To corroborate this classification, the surface expression level of each variant was statistically evaluated against the WT(1:1) control (a fixed theoretical value of 100%) using a two-tailed one-sample *t*-test, and against the WT(1:2) and WT(1:4) controls using a two-tailed unpaired *t*-test with Welch’s correction. Importantly, the qualitative trends of the *P*-values obtained from these tests corresponded well with the mean-based stratification, further supporting the validity of this cut-off.

Following this stratification, statistical differences in signaling responses (LogRAi) were assessed within each respective group. For variants in the WT(1:1)-matched group, a two-tailed one-sample *t*-test was performed against a theoretical baseline of 0. For variants in the WT(1:2)- matched group, a two-tailed unpaired *t*-test with Welch’s correction was performed against the WT(1:2) control. The R129^3.50^A mutant was excluded from this statistical analysis because its signaling response was completely abolished and yielded no measurable values. Because each point mutation was designed to independently evaluate the specific functional role of an individual residue, these analyses were treated as *a priori* planned comparisons. Therefore, unadjusted *P*-values were utilized to determine statistical significance without applying multiple comparison corrections, thereby maintaining analytical sensitivity. Statistical significance was defined as *P* < 0.05.

### Summary of MD simulations

We performed 16 independent simulations of the CX_3_CL1.44-US28-G_q_ complex starting from the T2C-state. All simulation lengths are shown in Supplementary Fig. 6. Simulation details are summarized in Supplementary Table 2.

### MD simulation setup

As the starting point for the simulations, we used a model of CX_3_CL1.44-US28-G_q_ in the T2C-state with certain unresolved regions modeled in using Coot. This model includes residues 17–306 of US28, residues 1–68 of CX_3_CL1.44, residues 1–359 of Gα_qiN18_, residues 3–340 of Gβ, and residues 7–68 of Gγ. The scFv was removed. The resulting model was aligned using the PPM 3 web server^55^.

The model was prepared for simulation using the Bilayer Builder in CHARMM-GUI^56–58^. Protein N- and C-termini were modeled as follows: GLYP (glycine N-terminus) and CTER (standard C-terminus) on Gα; ACE (acetylated N-terminus) and CTER on Gβ; ACE and CTER on Gγ; NTER (standard N-terminus) and CT3 (methylamidated C-terminus) on CX_3_CL1.44; and ACE and CT3 on US28. All titratable residues were kept in their standard protonation states at pH 7.4, except for the following US28 residues: Glu124^3.45^, Glu191^5.41^, Asp79^2.50^, and Asp128^3.49^, which were protonated (neutral). Glu124^3.45^ and Glu191^5.41^ were modeled as protonated because they are in TM regions that face the lipid membrane, where a negative charge is expected to be energetically unfavorable. Asp79^2.50^, and Asp128^3.49^ were modeled as neutral because there is evidence suggesting that these residues are protonated upon receptor activation^59,60^. All histidine residues were modeled as neutral with a proton on the epsilon nitrogen. Lipidations on the G protein were made as follows: GLYM (myristoylation) on Gly1 of Gα; CYSP (palmitoylation) on Cys2 of Gα; and CYSG (geranylgeranylation) on Cys68 of Gγ. Disulfide bridges were modeled between Cys8 and Cys34 of CX_3_CL1.44, between Cys12 and Cys50 of CX_3_CL1.44, between Cys23 and Cys263^7.25^ of US28, and between Cys104^3.25^ and Cys173^45.50^ of US28. The system was then embedded in a palmitoyloleoyl-phosphatidylcholine (POPC) bilayer with target dimensions of 130 Å × 130 Å and solvated using a water thickness of 20 Å with 150 mM neutralizing KCl. OpenMM input files with hydrogen-mass repartitioning (HMR) were produced at 310.15 K with the well-validated CHARMM36m forcefield parameters, including associated CHARMM36 lipid parameters, TIP3P water, as well as WYF parameters^61–64^. The prepared system contains 309,986 atoms, including 76,959 water molecules and 453 lipid molecules. This prepared system is the starting structure shown in Fig. 3a and Supplementary Fig. 6 and is considered *t* = 0 (the initial frame) in the analyzed trajectories.

### MD simulation protocols

Simulations were run with OpenMM (version 8.3.1)^65^ at 310.15 K using Langevin dynamics with a friction coefficient of 1 ps⁻¹ for temperature control. Electrostatic interactions were treated using the particle-mesh Ewald (PME) method with a real-space cutoff of 1.2 nm and an Ewald error tolerance of 5 × 10⁻⁴. Lennard-Jones interactions were computed with the same cutoff using a force-switching scheme between 1.0 and 1.2 nm. All bonds involving hydrogen atoms were constrained. For NPT equilibration and production stages, pressure was controlled at 1 bar using the Monte Carlo barostat with attempts every 100 steps.

Equilibration followed the standard protocol from Bilayer Builder in CHARMM-GUI. After 5000 steps of energy minimization, six restrained equilibration stages (stages 1–6) were performed. Stages 1–2 were run in the NVT ensemble for 0.125 ns per stage with a 1-fs timestep. Stage 3 was run in the NPT ensemble for 0.125 ns with a 1-fs timestep. Stages 4–6 were run in the NPT ensemble for 0.5 ns per stage with a 2-fs timestep, for a total of 1.875 ns of restrained equilibration after minimization. During equilibration, harmonic restraints were applied to protein and GDP during stages 1–6 and to lipids during stages 1–5. During stages 1–6, the restraint force constants were: 4000, 2000, 1000, 500, 200, 50 kJ·mol⁻¹·nm⁻² for protein backbone and GDP heavy-atom positions; 2000, 1000, 500, 200, 50, 0 kJ·mol⁻¹·nm⁻² for protein side-chain positions; 1000, 400, 400, 200, 40, 0 kJ·mol⁻¹·nm⁻² for lipid positions; and 1000, 400, 200, 200, 100, 0 kJ·mol⁻¹·rad⁻² for lipid dihedrals. After equilibration, production simulations were carried out in the NPT ensemble with a 4-fs timestep with coordinates written every 0.5 ns. No restraints were applied during production. Because these unbiased, conventional simulations revealed relevant transitions on the simulated timescales, we did not employ enhanced-sampling methods.

### MD simulation analysis protocols

Trajectories for analysis were concatenated and reimaged using catdcd-4.0b and MDTraj (version 1.11.0)^66^ at 10 ns per frame. Frames from equilibration stages 1–6 were not included in the analyzed trajectories. VMD (version 1.9.4a57)^67^ and MDTraj (version 1.11.0) were used for visual and quantitative analysis. For visual and quantitative analysis, the initial prepared structure was prepended as the first frame of each trajectory.

We monitored helicity in the membrane-distal segment of α5 by measuring the distance between the backbone oxygen atom of Asn336 and the backbone nitrogen atom of Val340. To measure AHD separation from the RHD, we used the distance between the Cα atoms of Asp146 and Tyr285. As the membrane-distal segment of α5 is helical both in the E- and C-states, but not in the T2C-state, we included another metric—the distance between the Cα atoms of Phe341 and Gln58 of Gα—to distinguish between an α5 that is translated “upward” (toward the membrane; as in the C-state) relative to the RHD and one that is translated “downward” (away from the membrane; as in the E- and inactive states) relative to the RHD. Based on these metrics and visual inspection, we consider simulations 1, 3, 7, 9, 14, and 16 (Supplementary Fig. 6) to exhibit partial transitions toward the C-state, and simulation 4 to exhibit a partial transition toward the E-state. The rest of the simulations—simulations 2, 5, 6, 8, 10, 11, 12, 13, and 15—generally retain characteristics of the T2C-state, and we thus did not consider them as exhibiting transitions (even partial ones) toward the C- or E-states. We note that this classification of trajectories into 3 strict categories involves some judgment; for instance, simulation 8 retains a broken membrane-distal α5 segment characteristic of the T2C-state, but the TCAT motif moves away from GDP in a manner unlike the remaining simulations in this category.

## Data availability

The cryo-EM maps and model coordinates have been deposited in the Electron Microscopy Data Bank (EMDB) and the Protein Data Bank (PDB) under accession codes EMD-66864/66942 (PDB: 9XH2/9XJP), EMD-66923 (PDB: 9XIY), EMD-66922/66928 (PDB: 9XIX/9XJ5), and EMD-66930 (PDB: 9XJ7) for the E-state, T2C-state, C-state, and C’-state complexes, respectively. Curated raw cryo-EM movies have been deposited in the Electron Microscopy Public Image Archive (EMPIAR) under accession code EMPIAR-13238. Simulation data have been deposited in a Zenodo repository under entry 20598927.

## Acknowledgments

We are grateful to Dr. Brian K. Kobilka for his guidance and helpful discussions. We thank Mses. Kayo Sato, Shigeko Nakano, and Ayumi Inoue (Tohoku University) for their assistance in plasmid construction and the NanoBiT assay, and Ms. Miho Sasaki (Cellular and Structural Physiology Laboratory; CeSPL) for research support. We also acknowledge Drs. John S. Burg and Timothy F. Miles for their contributions to earlier works, and Dr. Emily Weed-Nichols for their technical assistance with MD simulations. Cryo-EM data were collected at the Stanford cryo-EM center (cEMc). Google Gemini (3.1 Pro) was used to revise the text and enhance readability. The authors maintained full oversight, verified all generated content, and take full responsibility for the final manuscript.

## Contributions

K.M.J.: Data Curation (equal), Formal Analysis (equal), Investigation (equal), Methodology (supporting), Validation (equal), Visualization (equal), Writing – Original Draft (supporting), Writing – Review & Editing (equal). C.-M.S.: Data Curation (equal), Formal Analysis (lead), Investigation (supporting), Methodology (equal), Supervision (supporting), Validation (supporting), Visualization (supporting), Writing – Original Draft (supporting), Writing – Review & Editing (supporting). D.W., S.M.: Investigation (supporting), Writing – Review & Editing (supporting). Y.F.: Resources (supporting), Writing – Review & Editing (supporting). A.I.: Data Curation (supporting), Formal Analysis (supporting), Funding Acquisition (equal), Investigation (equal), Methodology (supporting), Resources (supporting), Supervision (supporting), Validation (supporting), Visualization (supporting), Writing – Original Draft (supporting), Writing – Review & Editing (supporting). K.C.G.: Conceptualization (equal), Funding Acquisition (lead), Methodology (equal), Resources (lead), Supervision (equal), Writing – Review & Editing (supporting). N.T.: Conceptualization (equal), Data Curation (equal), Formal Analysis (equal), Funding Acquisition (equal), Investigation (lead), Methodology (equal), Project Administration (lead), Supervision (equal), Validation (equal), Visualization (equal), Writing – Original Draft (lead), Writing – Review & Editing (equal).

## References

1. Alexander, S. P. H. et al. The Concise Guide to PHARMACOLOGY 2023/24: G protein-coupled receptors. Br. J. Pharmacol. 180 Suppl 2, S23–S144 (2023).

2. Zhang, M. et al. G protein-coupled receptors (GPCRs): advances in structures, mechanisms, and drug discovery. Signal Transduct. Target. Ther. 9, 88 (2024).

3. Conflitti, P. et al. Functional dynamics of G protein-coupled receptors reveal new routes for drug discovery. Nat. Rev. Drug Discov. 24, 251–275 (2025).

4. Hilger, D., Masureel, M. & Kobilka, B. K. Structure and dynamics of GPCR signaling complexes. Nat. Struct. Mol. Biol. 25, 4–12 (2018).

5. Weis, W. I. & Kobilka, B. K. The molecular basis of G protein-coupled receptor activation. Annu. Rev. Biochem. 87, 897–919 (2018).

6. Traut, T. W. Physiological concentrations of purines and pyrimidines. Mol. Cell. Biochem. 140, 1–22 (1994).

7. Dorsam, R. T. & Gutkind, J. S. G-protein-coupled receptors and cancer. Nat. Rev. Cancer 7, 79–94 (2007).

8. Jiang, H., Galtes, D., Wang, J. & Rockman, H. A. G protein-coupled receptor signaling: transducers and effectors. Am. J. Physiol. Cell Physiol. 323, C731–C748 (2022).

9. Herrera, L. P. T. et al. GPCRdb in 2025: adding odorant receptors, data mapper, structure similarity search and models of physiological ligand complexes. Nucleic Acids Res. 53, D425–D435 (2025).

10. Gusach, A. et al. Beyond structure: emerging approaches to study GPCR dynamics. Curr. Opin. Struct. Biol. 63, 18–25 (2020).

11. Papasergi-Scott, M. M. et al. Time-resolved cryo-EM of G-protein activation by a GPCR. Nature 629, 1182–1191 (2024).

12. Liu, X. et al. Structural insights into the process of GPCR-G protein complex formation. Cell 177, 1243–1251.e12 (2019).

13. Tsutsumi, N. et al. Atypical structural snapshots of human cytomegalovirus GPCR interactions with host G proteins. Sci. Adv. 8, eabl5442 (2022).

14. Dror, R. O. et al. SIGNAL TRANSDUCTION. Structural basis for nucleotide exchange in heterotrimeric G proteins. Science 348, 1361–1365 (2015).

15. Sun, X., Singh, S., Blumer, K. J. & Bowman, G. R. Simulation of spontaneous G protein activation reveals a new intermediate driving GDP unbinding. Elife 7, (2018).

16. Batebi, H. et al. Mechanistic insights into G-protein coupling with an agonist-bound G-protein-coupled receptor. Nat. Struct. Mol. Biol. 31, 1692–1701 (2024).

17. Robertson, M. J., et al. Non-equilibrium snapshots of ligand efficacy at the μ-opioid receptor. bioRxivorg (2025) doi:10.1101/2025.05.26.656223.

18. Maussang, D. et al. Human cytomegalovirus-encoded chemokine receptor US28 promotes tumorigenesis. Proc. Natl. Acad. Sci. U. S. A. 103, 13068–13073 (2006).

19. Humby, M. S. & O’Connor, C. M. Human Cytomegalovirus US28 is important for latent infection of hematopoietic progenitor cells. J. Virol. 90, 2959–2970 (2015).

20. Krishna, B. A. et al. Latency-associated expression of human Cytomegalovirus US28 attenuates cell signaling pathways to maintain latent infection. MBio 8, (2017).

21. Rosenkilde, M. M., Tsutsumi, N., Knerr, J. M., Kildedal, D. F. & Garcia, K. C. Viral G protein-coupled receptors encoded by β- and γ-herpesviruses. Annu. Rev. Virol. 9, 329–351 (2022).

22. Tsutsumi, N. et al. Insight into structural properties of viral G protein-coupled receptors and their role in the viral infection: IUPHAR Review 41. Br. J. Pharmacol. 182, 26–51 (2025).

23. Burg, J. S. et al. Structural biology. Structural basis for chemokine recognition and activation of a viral G protein-coupled receptor. Science 347, 1113–1117 (2015).

24. Miles, T. F. et al. Viral GPCR US28 can signal in response to chemokine agonists of nearly unlimited structural degeneracy. Elife 7, (2018).

25. Huang, S. K. et al. Mapping the conformational landscape of the stimulatory heterotrimeric G protein. Nat. Struct. Mol. Biol. 30, 502–511 (2023).

26. Gregorio, G. G. et al. Single-molecule analysis of ligand efficacy in β2AR-G-protein activation. Nature 547, 68–73 (2017).

27. Maeda, S. et al. Development of an antibody fragment that stabilizes GPCR/G-protein complexes. Nat. Commun. 9, 3712 (2018).

28. Zhang, D. et al. Structural insights into angiotensin receptor signaling modulation by balanced and biased agonists. EMBO J. 42, e112940 (2023).

29. Lu, M. et al. Activation of the human chemokine receptor CX3CR1 regulated by cholesterol. Sci. Adv. 8, eabn8048 (2022).

30. Zhang, H. et al. Structural basis for chemokine recognition and receptor activation of chemokine receptor CCR5. Nat. Commun. 12, 4151 (2021).

31. Heo, Y. et al. Structure of the human galanin receptor 2 bound to galanin and Gq reveals the basis of ligand specificity and how binding affects the G-protein interface. PLoS Biol. 20, e3001714 (2022).

32. Fraile-Ramos, A. et al. The human cytomegalovirus US28 protein is located in endocytic vesicles and undergoes constitutive endocytosis and recycling. Mol. Biol. Cell 12, 1737–1749 (2001).

33. Waldhoer, M. et al. The carboxyl terminus of human cytomegalovirus-encoded 7 transmembrane receptor US28 camouflages agonism by mediating constitutive endocytosis. J. Biol. Chem. 278, 19473–19482 (2003).

34. Zhou, Q. et al. Common activation mechanism of class A GPCRs. Elife 8, (2019).

35. Nishimura, A. et al. Structural basis for the specific inhibition of heterotrimeric Gq protein by a small molecule. Proc. Natl. Acad. Sci. U. S. A. 107, 13666–13671 (2010).

36. Hu, Q. & Shokat, K. M. Disease-causing mutations in the G protein Gαs subvert the roles of GDP and GTP. Cell 173, 1254–1264.e11 (2018).

37. Takasaki, J. et al. A novel Galphaq/11-selective inhibitor. J. Biol. Chem. 279, 47438–47445 (2004).

38. Mühle, J. et al. Cyclic peptide inhibitors function as molecular glues to stabilize Gq/11 heterotrimers. Proc. Natl. Acad. Sci. U. S. A. 122, e2418398122 (2025).

39. Pándy-Szekeres, G. et al. The G protein database, GproteinDb. Nucleic Acids Res. 50, D518–D525 (2022).

40. Khan, S. et al. Structural snapshots capture nucleotide release at the μ-opioid receptor. Nature (2025) doi:10.1038/s41586-025-09677-6.

41. Tsutsumi, N. et al. Structural basis for the constitutive activity and immunomodulatory properties of the Epstein-Barr virus-encoded G protein-coupled receptor BILF1. Immunity 54, 1405–1416.e7 (2021).

42. Schorb, M., Haberbosch, I., Hagen, W. J. H., Schwab, Y. & Mastronarde, D. N. Software tools for automated transmission electron microscopy. Nat. Methods 16, 471–477 (2019).

43. Punjani, A., Rubinstein, J. L., Fleet, D. J. & Brubaker, M. A. cryoSPARC: algorithms for rapid unsupervised cryo-EM structure determination. Nat. Methods 14, 290–296 (2017).

44. Bepler, T. et al. Positive-unlabeled convolutional neural networks for particle picking in cryo-electron micrographs. Nat. Methods 16, 1153–1160 (2019).

45. Pettersen, E. F. et al. UCSF ChimeraX: Structure visualization for researchers, educators, and developers. Protein Sci. 30, 70–82 (2021).

46. Liebschner, D. et al. Macromolecular structure determination using X-rays, neutrons and electrons: recent developments in Phenix. Acta Crystallogr. D Struct. Biol. 75, 861–877 (2019).

47. Abramson, J. et al. Accurate structure prediction of biomolecular interactions with AlphaFold 3. Nature 630, 493–500 (2024).

48. Read, R. J., Millán, C., McCoy, A. J. & Terwilliger, T. C. Likelihood-based signal and noise analysis for docking of models into cryo-EM maps. Acta Crystallogr. D Struct. Biol. 79, 271–280 (2023).

49. Millán, C., McCoy, A. J., Terwilliger, T. C. & Read, R. J. Likelihood-based docking of models into cryo-EM maps. Acta Crystallogr. D Struct. Biol. 79, 281–289 (2023).

50. Emsley, P., Lohkamp, B., Scott, W. G. & Cowtan, K. Features and development of coot. Acta Crystallogr. D Biol. Crystallogr. 66, 486–501 (2010).

51. Kabsch, W. & Sander, C. Dictionary of protein secondary structure: pattern recognition of hydrogen-bonded and geometrical features. Biopolymers 22, 2577–2637 (1983).

52. Chen, V. B. et al. MolProbity: all-atom structure validation for macromolecular crystallography. Acta Crystallogr. D Biol. Crystallogr. 66, 12–21 (2010).

53. Morin, A. et al. Collaboration gets the most out of software. Elife 2, e01456 (2013).

54. Niwa, H., Yamamura, K. & Miyazaki, J. Efficient selection for high-expression transfectants with a novel eukaryotic vector. Gene 108, 193–199 (1991).

55. Lomize, A. L., Todd, S. C. & Pogozheva, I. D. Spatial arrangement of proteins in planar and curved membranes by PPM 3.0. Protein Sci. 31, 209–220 (2022).

56. Gao, Y. et al. CHARMM-GUI supports hydrogen mass repartitioning and different protonation states of phosphates in lipopolysaccharides. J. Chem. Inf. Model. 61, 831–839 (2021).

57. Lee, J. et al. CHARMM-GUI input generator for NAMD, GROMACS, AMBER, OpenMM, and CHARMM/OpenMM simulations using the CHARMM36 additive force field. J. Chem. Theory Comput. 12, 405–413 (2016).

58. Jo, S., Kim, T., Iyer, V. G. & Im, W. CHARMM-GUI: a web-based graphical user interface for CHARMM. J. Comput. Chem. 29, 1859–1865 (2008).

59. Ghanouni, P. et al. The effect of pH on beta(2) adrenoceptor function. Evidence for protonation-dependent activation. J. Biol. Chem. 275, 3121–3127 (2000).

60. Ranganathan, A., Dror, R. O. & Carlsson, J. Insights into the role of Asp79(2.50) in β2 adrenergic receptor activation from molecular dynamics simulations. Biochemistry 53, 7283–7296 (2014).

61. Huang, J. et al. CHARMM36m: an improved force field for folded and intrinsically disordered proteins. Nat. Methods 14, 71–73 (2017).

62. Klauda, J. B. et al. Update of the CHARMM all-atom additive force field for lipids: validation on six lipid types. J. Phys. Chem. B 114, 7830–7843 (2010).

63. Jorgensen, W. L., Chandrasekhar, J., Madura, J. D., Impey, R. W. & Klein, M. L. Comparison of simple potential functions for simulating liquid water. J. Chem. Phys. 79, 926–935 (1983).

64. Khan, H. M., MacKerell, A. D., Jr & Reuter, N. Cation-π interactions between methylated ammonium groups and tryptophan in the CHARMM36 additive force field. J. Chem. Theory Comput. 15, 7–12 (2019).

65. Eastman, P. et al. OpenMM 8: Molecular dynamics simulation with machine learning potentials. J. Phys. Chem. B 128, 109–116 (2024).

66. McGibbon, R. T. et al. MDTraj: A modern open library for the analysis of molecular dynamics trajectories. Biophys. J. 109, 1528–1532 (2015).

67. Humphrey, W., Dalke, A. & Schulten, K. VMD: visual molecular dynamics. J. Mol. Graph. 14, 33–8, 27–8 (1996).

