## Supplementary Information for "Structural basis of stepwise G protein activation by the viral chemokine receptor US28"

Correspondence:

### Supplementary Discussion

#### General 3D classification strategy and interpretation of the C-state variants

In the $CX_3CL1.44$-US28-$G_q$ structures, the globular body of the $CX_3CL1.44$ chemokine exhibited flexibility (Supplementary Fig. 2), a feature commonly observed in the wild-type $CX_3CL1$-US28 complex[1] and human chemokine receptor systems[2,3]. In contrast, the engineered N-terminal seven residues of the chemokine were well-defined. Therefore, to maximize the resolution of the US28-$G_q$ interface, our primary 3D refinements were performed with all monomeric particles, regardless of heterogeneity in the chemokine density. When a subclass of particles showed the chemokine body in a metastable conformation, we also generated a separate reconstruction to visualize its position (Supplementary Fig. 2,3).

The C-state US28-$G_q$ interface also exhibited flexibility, where we focused on the particle subset that yielded the highest-resolution features. The stochastic motion suggests that US28 can move further away from the Gβ interface, which leads to a more exclusive interaction with the extended α5 helix, and may represent a mechanism to ensure swift dissociation of the complex upon GTP binding.

Among the C-state variants, the class with closed AHD, termed the C'-state, was also identified from the 3D classification that identified the T2C-state, and refined to 2.9 Å resolution. Thus, it resembles the T2C-state with a closed AHD but lacks a clear GDP density (Supplementary Fig. 5c). Given its higher local conformational similarity to the C-state around the nucleotide-binding site, it most likely represents a state arising from stochastic AHD closure after C-state formation[4], although it could alternatively represent an intermediate between the T2C- and C-states. For the T2C- and C'-states, key regions were locally refined to guide model building (Supplementary Fig. 2).

#### Detailed structural analysis of the US28-$G_q$ interfaces

##### *State-dependent interactions at the $Gα_q$-αN helix*

The engagement of US28 with the $Gα_q$-αN helix shifts dynamically during the activation trajectory (Fig. 4a and Supplementary Fig. 7). In the E-state, US28 recognizes a more N-terminal region of αN. Specifically, the $Gα_q$-Arg30 sidechain contacts the US28-Val141 sidechain and US28-

Gln65 Cα, while the Gα$_q$-Arg34 sidechain forms a hydrogen bond with the US28-Tyr138 backbone carbonyl and the Gα$_q$-Arg37 sidechain interacts with the Tyr138 sidechain. As the complex transitions, the interface shifts to a more C-terminal region in the subsequent states: in the T2C-state, the Gα$_q$-Arg37 guanidinium group shifts slightly to stack on the face of US28-Tyr138 while Gα$_q$-Arg38 engages the edge of US28-Tyr138, and in the C-state, Gα$_q$-Arg38 additionally forms a hydrogen bond with the US28-Met136 backbone carbonyl.

This interaction mode in the C-state complex is distinct from that of typical human chemokine receptors. Due to its shorter ICL2, US28 sits more shallowly within the Gα$_q$ crevice (formed by the α5 helix, αN-β1 junction, and β2-β3 loop) and therefore does not engage it as deeply. This distinction is important because the αN-β1 junction is one of the major sensors for receptor activation, and the connected β1-β3 sheet serves as a communication hub to activate the G protein[5].

#### *Detailed rearrangements at the Gα$_q$-α5 helix*

The most substantial interface remodeling occurs at the Gα$_q$ C-terminus, driven by the formation and rotation of the α5 helix (Fig. 4a and Supplementary Fig. 7). This movement pivots around the Gα$_q$-Val(-1) (where -1 is the extreme C-terminus; preceding positions are -2, -3, etc.) residue, which serves as a conserved anchor point. This C-terminal anchor is stabilized by the interaction of the carboxylate with the N-terminal dipole of US28 H8, which unwinds slightly in the T2C and C states to donate hydrogen bonds from the backbone and sidechain of US28-Thr296. The interactions of the preceding Asn(-3) residue, however, are state-dependent. In the E-state, its sidechain forms a single hydrogen bond with the US28-Arg129 sidechain. In contrast, in the C-state, it forms an extensive hydrogen bonding network with the backbones of TM7 and the TM7-H8 turn. The Tyr(-4) sidechain also undergoes a marked rearrangement of its interactions. In the E-state, its hydroxyl group forms hydrogen bonds with US28-Ser67 and -Gly68. In the T2C- and C-states, however, the sidechain reorients such that its aromatic ring is sandwiched between US28-Cys66/Arg129 and is ultimately stabilized in the C-state by a thiol-π interaction, while the positive quadrupole of the ring edge is oriented toward the side chain carboxylic acid of US28-Asp128. The engagement at the base of the intracellular pocket is also state-dependent. In the E-state, the Leu(-9) residue is lodged at the pocket's edge on ICL3, anchored by both a backbone-backbone hydrogen bond and sidechain hydrophobic interactions with US28-Q219. In the T2C- and C-states, however, the interface rearranges, allowing the Asp(-14) residue to insert deeper into this pocket (Fig. 4a). Consequently, the formation of the α-helix draws a C-terminal segment of $G\alpha_q$, previously positioned outside the pocket in the E-state, into the intracellular cleft. This engagement facilitates the formation of numerous new hydrophobic and polar interactions (Supplementary Fig. 7).

***Comparison between the C-state US28 complexes with $G_q$ and $G_i$***

Interestingly, the previously identified C-state US28-$G_i$ structure retains a US28-mediated bridge between $G\alpha_i$ and Gβ, a feature attributed to a unique bend in its α5 helix[1]. We propose that the relaxation of this bending strain would straighten the α5 helix and cause a slight rotational displacement of the G protein (Fig. 3d), resulting in the predominantly US28-Gα interface observed in the current $G_q$ complex. Therefore, the newly resolved C-state $CX_3CL1.44$-US28-$G_q$ complex represents a more canonical and fully active conformation compared to that of the US28-$G_i$ complex. This structural difference likely underlies a reduced GEF activity of US28 toward $G_i$ compared to $G_{q/11}$ (ref [1]).

**Structural determinants in G protein subtype selectivity by US28**

Our structures of the signaling-competent, authentic C-state US28-$G_q$ complex provide a structural framework to understand the G protein subtype selectivity of US28, which is known to couple to $G_q$, $G_{12/13}$, and $G_i$, but not $G_s$[6,7]. A comparison of the C-terminal sequences of Gα subunits, which primarily determine the coupling specificity, shows a key difference, where $G\alpha_s$ uniquely possesses a glutamate residue at the -3 position (Supplementary Fig. 10a). To visualize the structural consequence of this residue, we generated a homology model of the US28-$G\alpha_s$ complex based on the C-state US28-$G_q$ structure[8].

The homology model predicts significant electrostatic repulsion between the negatively charged sidechain of $G\alpha_s$-Glu(-3) and the backbone carbonyls at the intracellular end of US28 TM7. This contrasts with the favorable interactions observed in the C-state US28-$G_q$ complex and the accommodation of the smallest sidechain in the US28-$G_i$ complex (Supplementary Fig. 10b,c). While $G\alpha_s$ could be compatible with the E-state by interacting with US28-Arg129, this clash in the C-state would likely prevent the stable engagement required for G protein activation.

This predicted inhibitory mechanism in US28 contrasts with the established mode of $G_s$ coupling by prototypical $G_s$-coupled receptors, such as β2AR[9] (Supplementary Fig. 10b,c). In the active β2AR-$G_s$ complex structure, TM6 undergoes a much larger outward displacement compared to that of US28. This wider opening of the intracellular cavity repositions the $G\alpha_s$ C-terminus, placing the Glu(-3) residue at the periphery of the binding interface rather than at its core. In this conformation, the Glu(-3) sidechain makes minimal direct contact with the receptor and is instead accommodated by weak interactions between TM6 and TM7. Therefore, the more compact intracellular architecture of US28 provides a structural basis for its inability to couple with $G_s$, offering a clear mechanism for its G protein selectivity.

This structural constraint imposed by a relatively narrow intracellular opening may also explain the G protein coupling profiles of endogenous human chemokine receptors[10,11]. These receptors predominantly signal through $G_{i/o}$, which has the least bulky C-terminal region, and show some coupling to $G_{q/11}$ and $G_{12/13}$ subtypes but generally exclude $G_s$[10].

### Structural basis for G protein-biased agonism by engineered chemokines

The $CX_3CL1.44$-US28 structure determined here allows us to compare the structures of US28 bound to three different $CX_3CL1$ variants, including the crystal structure of $CX_3CL1.35$-US28 (Supplementary Fig. 11a-d). Notably, the N-terminal peptides of the wild-type $CX_3CL1$ and its G protein-biased variants, $CX_3CL1.35$ and $CX_3CL1.44$, each trace a unique path at CRS2.

The N-terminal peptide of $CX_3CL1.44$ forms an α-helix, allowing it to project sidechains into both the major and minor ligand-binding pockets. The peptide of $CX_3CL1.35$, while lacking a defined secondary structure, also engages both pockets. In contrast, the unstructured N-terminal peptide of the wild-type $CX_3CL1$ interacts solely with the minor pocket. The interaction at the major pocket induces displacement of TM5 and TM6. Furthermore, a critical difference is observed at CRS3. The wild-type chemokine, which is endogenously β-arrestin-biased, establishes extensive contacts between the chemokine globular core and receptor extracellular loop (ECL) 2 at CRS3 (Supplementary Fig. 11e). This interaction, which appears to induce a downward conformational change in ECL2, is absent in the two G protein-biased engineered variants.

Therefore, we propose a two-part mechanism for the G protein-selective activation of US28: (1) the simultaneous engagement of both the major and minor binding pockets by the chemokine's N-terminal peptide, and (2) the absence of the specific CRS3-mediated interaction that repositions ECL2.

### Supplementary Table 1 | Cryo-EM data collection and refinement statistics

| | E-state | | T2C-state | | C'-state | | C-state | |
|---|---|---|---|---|---|---|---|---|
| | CX$_3$CL1 core unmodeled | CX$_3$CL1 core modeled | CX$_3$CL1 core unmodeled with GDP | | CX$_3$CL1 core unmodeled without GDP | | CX$_3$CL1 core unmodeled | CX$_3$CL1 core modeled |
| | global | global | global | local | global | local | global | global |
| **EMDB ID (EMD-)** | **66864** | **66942** | **66923** | N/A | **66930** | N/A | **66922** | **66928** |
| **PDB ID** | **9XH2** | **9XJP** | **9XIY** | N/A | **9XJ7** | N/A | **9XIX** | **9XJ5** |
| **Map** | 1 | 2 | 5-1 | 5-2/5-3 | 6 -1 | 6-2/6-3 | 3 | 4 |
| Nominal mag. | **105,000x** | | | | | | | |
| Calibrated mag. | **57,624x** | | | | | | | |
| Voltage (kV) | **300** | | | | | | | |
| Exposure(e/Å$^2$) | **50** | | | | | | | |
| Defocus range (μm) | **-0.8 to -2.0** | | | | | | | |
| Pixel size (Å) | **0.8677** | | | | | | | |
| Symmetry | **C1** | | | | | | | |
| Initial particle no. | **8,400,483** | | | | | | | |
| Final particle no. | 193,867 | 64,658 | 70,394 | US28/$G_q$ 69,802/ 69,799 | 69,523 | US28/$G_q$ 68,903/ 68,894 | 268,599 | 89,862 |
| Resolution (Å) FSC=0.143 (masked) | 2.6 | 2.7 | 2.9 | 3.5/3.2 | 2.9 | 3.4/3.2 | 2.6 | 2.8 |
| Sharpening *B* (Å$^2$)* | -78.3 | -68.1 | -40 | -88.5/-84.7 | -73.4 | -86.5/-81.3 | -86.4 | -75.3 |
| **Model** | | | | | | | | |
| Initial model | PDB: 7RKF, AlphaFold3 | PDB: 7RKF, AlphaFold3 | E-state model | N/A | T2C-state model | N/A | PDB: 7RKF, AlphaFold3 | PDB: 7RKF, AlphaFold3 |
| Resolution (Å) FSC = 0.5 (masked) | 2.8 | 2.9 | 3.2 | | 3.2 | | 2.8 | 3.0 |
| Model composition | | | | | | | | |
| Non-H atoms | 10,161 | 10,700 | 10,005 | | 10,068 | | 9,083 | 9,554 |
| Protein residues | 1,277 | 1,343 | 1,260 | | 1,270 | | 1,152 | 1,213 |
| Ligands | GDP/CLR | CLR | GDP | | N/A | | N/A | N/A |
| *B* factors (Å$^2$) | | | | | | | | |
| Protein | 48.0 | 44.2 | 110.79 | | 68.1 | | 55.5 | 58.6 |
| Ligand | 25.9/49.0 | 62.4 | 59.60 | | N/A | | N/A | N/A |
| R.m.s. deviations | | | | | | | | |
| Bond lengths (Å) | 0.003 | 0.003 | 0.002 | | 0.003 | | 0.003 | 0.003 |
| Bond angles (°) | 0.550 | 0.508 | 0.453 | | 0.544 | | 0.537 | 0.551 |
| Validation | | | | | | | | |
| MolProbity score | 1.68 | 1.48 | 1.58 | | 2.00 | | 1.41 | 1.74 |
| Clashscore | 3.76 | 3.43 | 4.02 | | 6.85 | | 3.16 | 4.16 |
| Rotamer outliers (%) | 3.54 | 2.77 | 2.39 | | 4.01 | | 2.42 | 3.46 |
| Ramachandran plot | | | | | | | | |
| Favored (%) | 97.55 | 97.97 | 97.50 | | 97.05 | | 97.97 | 97.32 |
| Allowed (%) | 2.45 | 2.03 | 2.50 | | 2.87 | | 2.03 | 2.68 |
| Disallowed (%) | 0 | 0 | 0 | | 0.08 | | 0 | 0 |
| EMRinger score | 3.14 | 3.57 | 2.18 | | 2.58 | | 3.47 | 3.21 |

*Uniformly sharpened with indicated *B* factors for Phenix refinement and/or local map visualization. DeepEMhancer sharpened for overall map representation.

**Supplementary Table 2 | Summary of simulation setup.**

| | |
|---|---|
| **initial dimensions** | 130 Å × 130 Å × 195 Å |
| **atoms** | 309,986 |
| **waters** | 76,959 |
| **lipids** | 453 POPC |
| **neutralizing salt** | 0.15 M KCl |

*System setup for all 16 simulations started from the T2C-state.

Simulation lengths are listed in Supplementary Fig 6.

### Supplementary Table 3 | Summary of structure-guided mutagenesis at the US28-$G_q$ interface.

| **Mutant** | State-dependence of contact | Structural hypothesis | Expression [%WT(1:1) ± SEM] | LogRAi [mean ± SEM] | *P*-value | Number of independent experiments |
|---|---|---|---|---|---|---|
| **R129$^{3.50}$A** | Persistent across all states | Primary anchor for G protein and essential for active US28 conformation | 73.3 ± 10.8 | ND (Completely abolished) | ND | 3 |
| **D128$^{3.49}$A** | Persistent across all states | Maintains interaction with Tyr(-4) of G$\alpha_q$ C-terminus throughout activation | 53.0 ± 2.9 | -0.64 ± 0.16 | 0.0236 [vs. WT(1:2)] | 4 |
| **S218$^{5.68}$A** | Persistent across all states | Provides continuous structural support | 79.8 ± 8.8 | -0.67 ± 0.02 | < 0.0001 [vs. WT(1:2)] | 3 |
| **V294$^{7.56}$A** | None (Allosteric) | Receptor stabilization via packing against V229$^{6.36}$ | 96.2 ± 9.5 | -0.64 ± 0.21 | 0.0897 [vs. WT(1:1)] | 3 |
| **K297$^{8.49}$A** | Forms in T2C- and C-states | Supports later activation steps | 76.3 ± 4.2 | -1.00 ± 0.05 | 0.0004 [vs. WT(1:2)] | 3 |
| **R221$^{ICL3}$A** | Greatest in E-state, decreases in T2C-/C-states | Supports initial engagement | 108.1 ± 9.5 | -0.20 ± 0.01 | 0.0044 [vs. WT(1:1)] | 3 |
| **S67$^{2.38}$A** | Extensive in E-/T2C-states, decreases in C-state | Supports early activation steps | 83.5 ± 2.9 | -0.33 ± 0.06 | 0.0092 [vs. WT(1:1)] | 4 |
| **I133$^{3.54}$A** | Transiently increased in T2C-state | Selectively supports T2C-state | 82.1 ± 7.0 | -0.84 ± 0.04 | 0.0002 [vs. WT(1:1)] | 4 |
| **R225$^{6.32}$A** | Transiently increased in T2C-state | Selectively supports T2C-state | 94.5 ± 4.2 | -0.28 ± 0.05 | 0.0258 [vs. WT(1:1)] | 3 |

ND: Not Determined.

Each independent experiment was performed in technical triplicate.

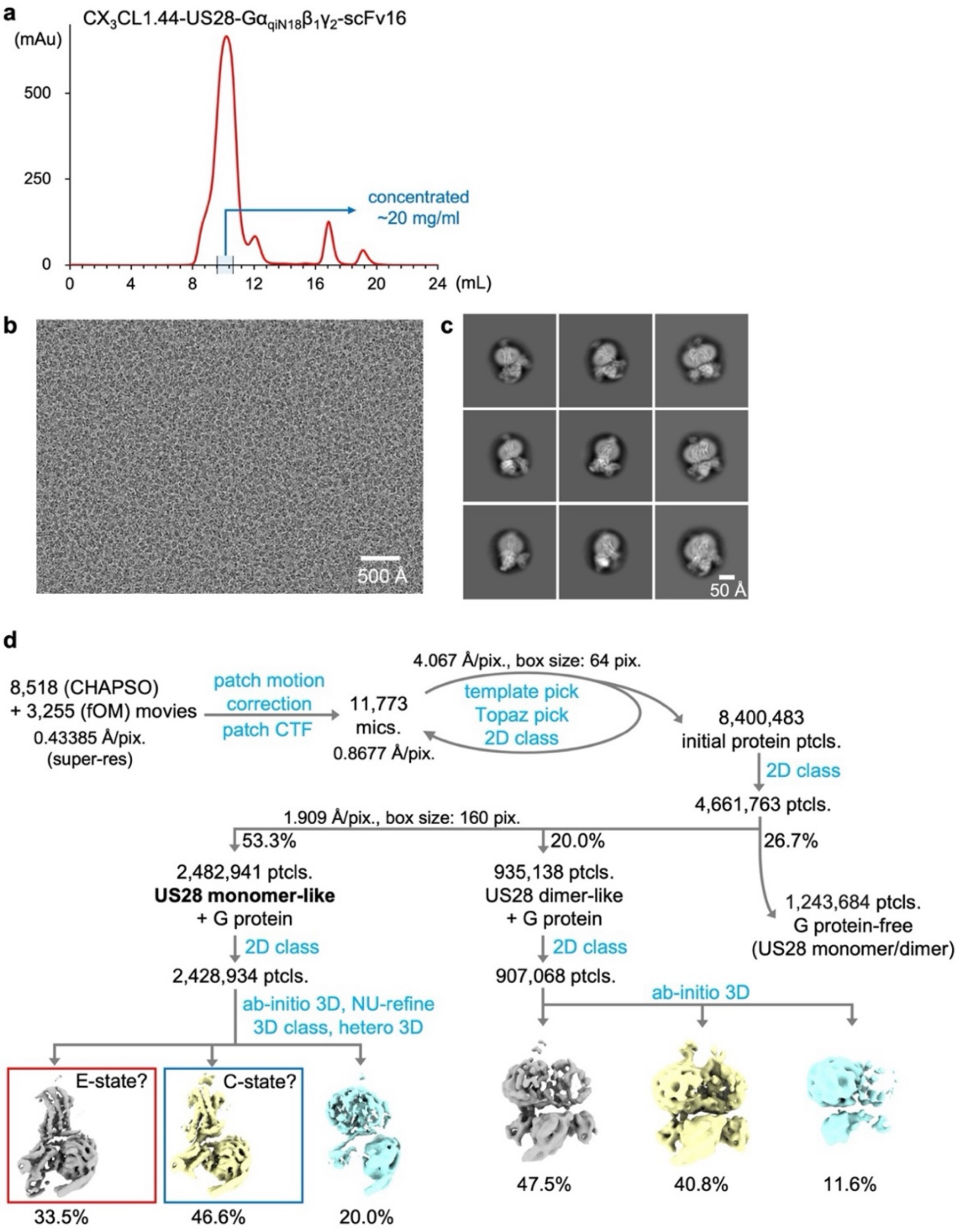


**Supplementary Fig. 1| Purification of the $CX_3CL1.44$-US28-$G_q$ complex and the initial cryo-EM data processing scheme.**

**a**, Size-exclusion chromatography (SEC) profile of the purified $CX_3CL1.44$-US28-$G\alpha_{qiN18}\beta_1\gamma_2$-scFv16 complex. **b**, Representative cryo-EM micrograph (scale bar 500 Å) and **c**, selected 2D class averages (left, scale bar 50 Å). A low-resolution class average from datasets collected in thicker ice areas is also shown (bottom right), suggesting a potential 2:2 receptor-$G_q$ complex in which two $CX_3CL1.44$ and two $G_q$ molecules are visualized. **d**, Flowchart of the initial cryo-EM data processing. Movies collected from samples prepared with CHAPSO and fOM were jointly processed. Initial 2D and 3D classifications separated the dataset into US28 monomer-like, US28 dimer-like, and G protein-free populations. The monomer-like population showed initial heterogeneity corresponding to putative E-state and C-state conformations. Patch CTF: Patch CTF Estimation, template pick: Template Picker, Topaz pick: training and extraction by Topaz, 2D class: 2D Classification, ab initio 3D: Ab-Initio Reconstruction, NU-refine: Non-uniform Refinement, 3D class: 3D Classification, hetero 3D: Heterogeneous Refinement.

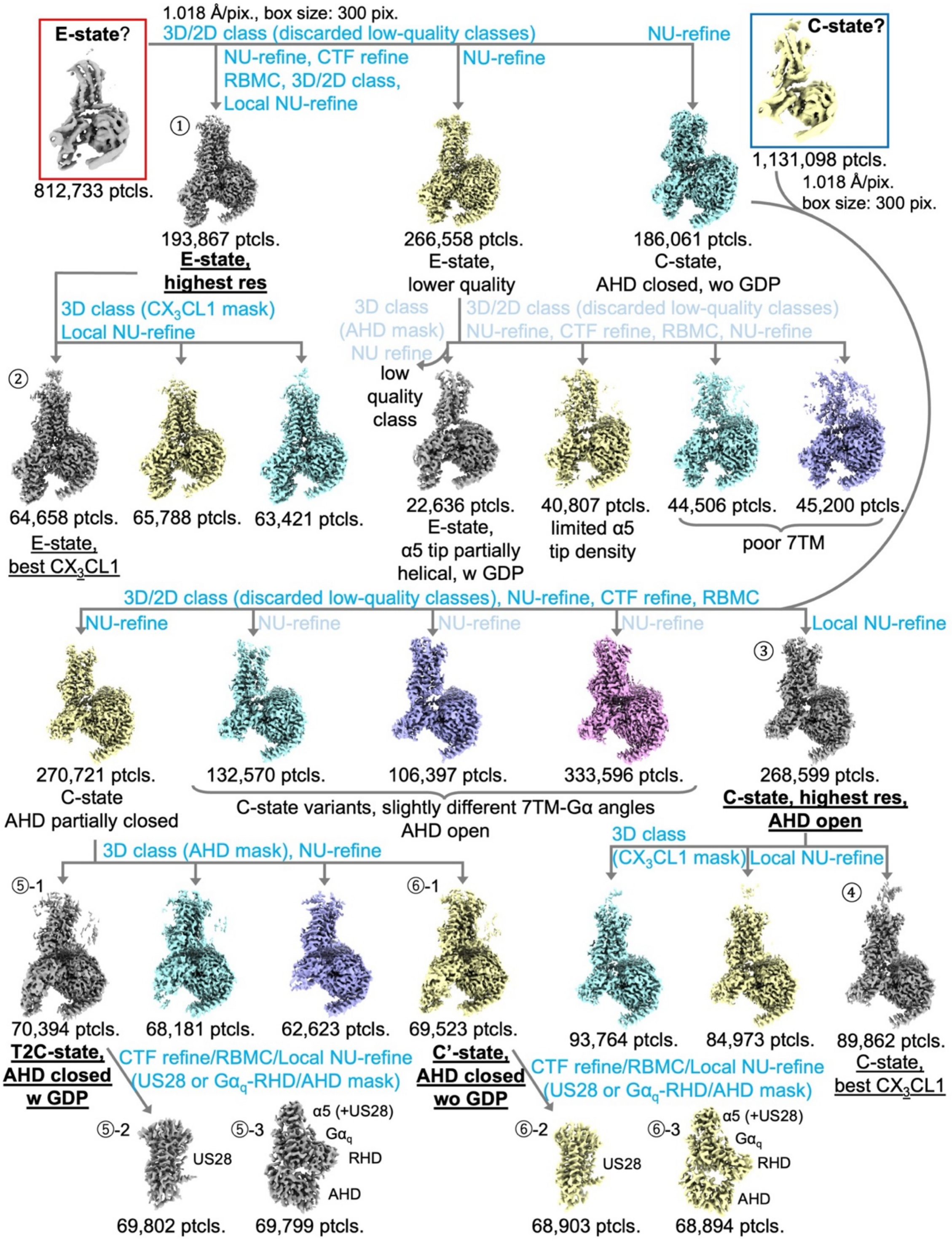


**Supplementary Fig. 2| Detailed 3D classification workflow for the monomeric US28-$G_q$ complex.**
Flowchart detailing the iterative classification and refinement strategy for the monomeric particles. Starting from the initial E-state-like and C-state-like populations, extensive processing using 3D/2D classification, (local) NU-refine, CTF refinement, and RBMC was performed. This pipeline resolved the high-resolution structures for the E-state (1, 2), C-state (3, 4), T2C-state (5-1), and C'-state (6-1). Mask-focused classifications (e.g., on $CX_3CL1$ body or AHD) and local refinements (5-2, 5-3, 6-2, 6-3) were employed to resolve heterogeneity and improve map quality in specific regions. RBMC: Reference-Based Motion Correction, CTF-refine: Global and Local CTF refinement, Local NU-refine: Local Refinement with NU-refine.

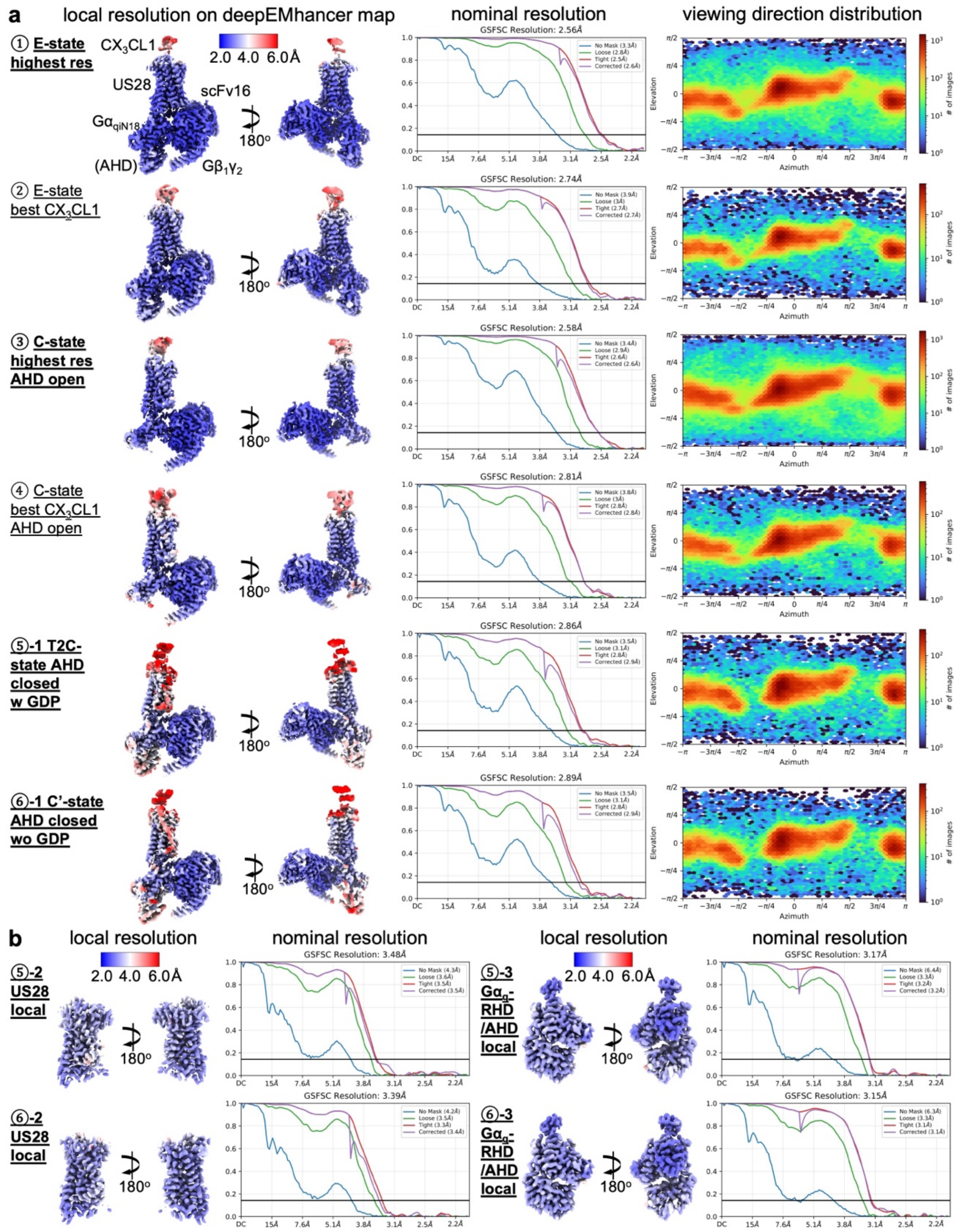


**Supplementary Fig. 3| Quality assessment of the cryo-EM reconstructions.**

**a**, Evaluation of the globally refined cryo-EM maps for the major conformational states: (1) E-state, highest resolution; (2) E-state, best $CX_3CL1$ core density; (3) C-state, highest resolution, open AHD; (4) C-state, best $CX_3CL1$ core density, open AHD; (5-1) T2C-state, closed AHD with GDP; and (6-1) C'-state, closed AHD without GDP. For each state, the local resolution map (colored on the deepEMhancer map), the gold-standard Fourier Shell Correlation (GSFSC) curve indicating the nominal resolution, and the viewing direction distribution plot are shown. **b**, Evaluation of the locally refined maps used as model building guides for the T2C-state (5-2: US28 local; 5-3: $G\alpha_q$-RHD/AHD local) and the C'-state (6-2: US28 local; 6-3: $G\alpha_q$-RHD/AHD local).

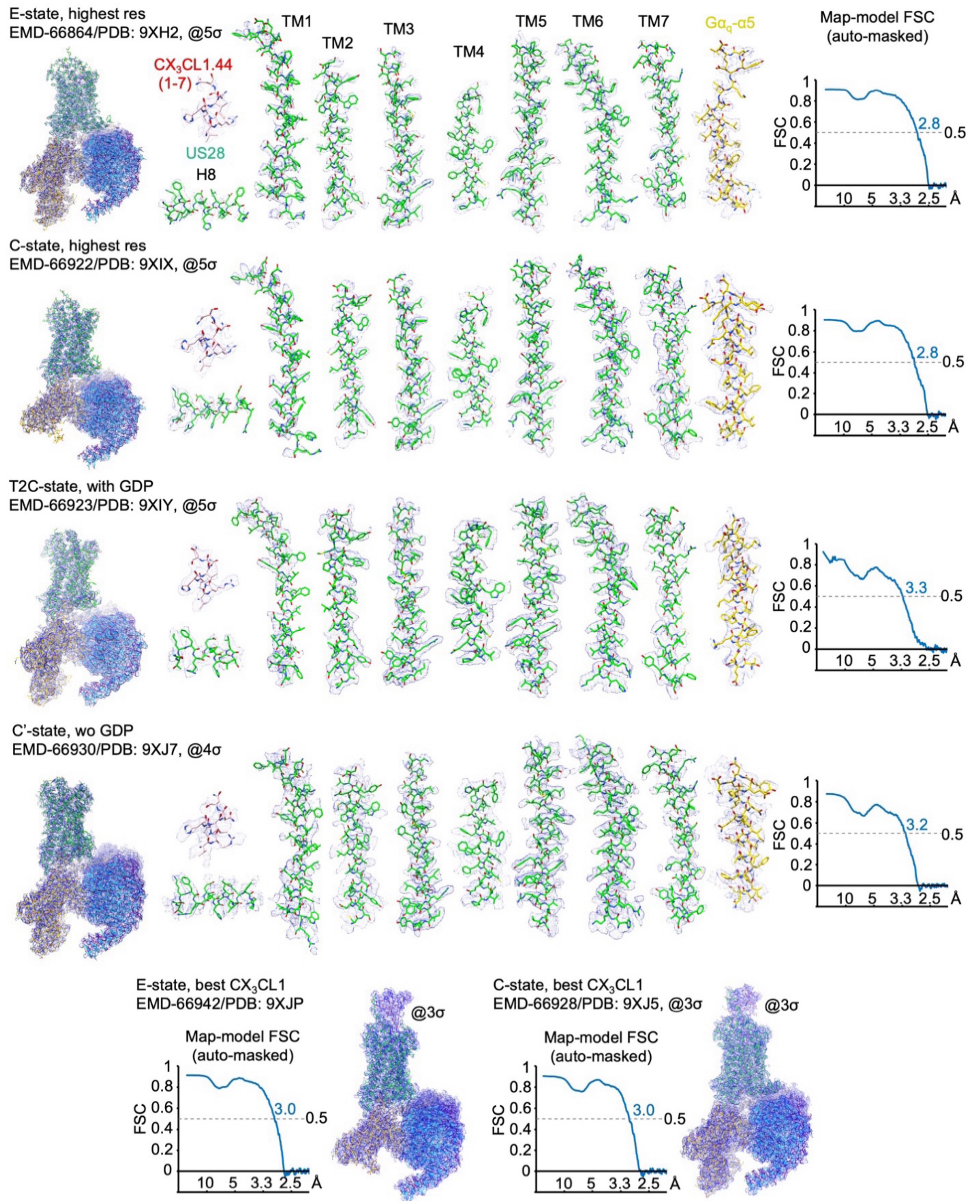


**Supplementary Fig. 4 | Representative cryo-EM densities and model validation.**
Representative cryo-EM densities (blue mesh) overlaid with the refined atomic models for the E-state, C-state, T2C-state, and C'-state. Densities are shown for key structural elements including the $CX_3CL1.44$ N-terminus (residues 1-7), US28 transmembrane helices (TMs) and helix 8 (H8), and the $G\alpha_q$-α5 helix. Maps are contoured at the indicated sigma (σ) levels. The map-versus-model Fourier Shell Correlation (FSC) curves (auto-masked) are shown for each state.

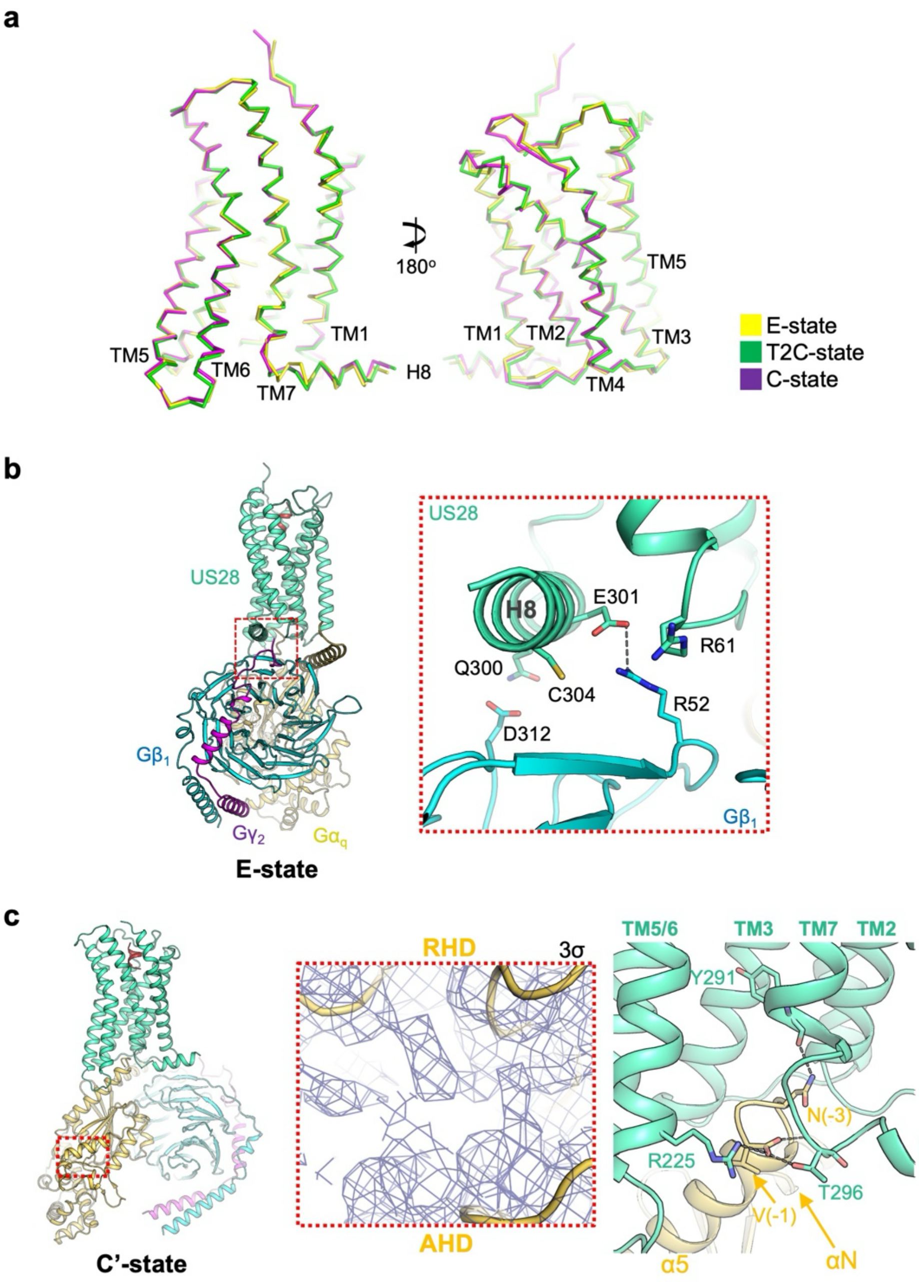


**Supplementary Fig. 5 | US28 conformation across distinct G protein engaging states, interface between US28 and Gβ in the E-state, and overall view of the C'-state.**

**a**, Structural alignment of US28 bound to $G_q$ across the E-state (yellow), T2C-state (green), and C-state (purple). The receptors are represented as ribbon models. The structures exhibit a high degree of similarity with a global Cα RMSD of ~0.6 Å, demonstrating that US28 acts as a rigid catalytic scaffold during progressive G protein engagement, while subtle state-dependent conformational changes beyond the resolution limits of the current static cryo-EM maps cannot be formally excluded. **b**, Interface between US28 and Gβ in the E-state. Overall (left) and close-up (right) views of the US28-Gβ interface in the E-state. Contacting residues are shown as sticks and a salt bridge is indicated by the gray dashed line. **c**, Views of the nucleotide-binding pocket and interface between the US28 intracellular core (green-cyan) and the $G\alpha_q$ C-terminus (yellow) of the $CX_3CL1.44$-US28-$G_q$ complex in the nucleotide-free C'-state. The panels show an overall view (left), a close-up of the nucleotide-binding pocket (middle), and the receptor-Gα C-terminus interface (right), in a layout analogous to Fig. 2b,d. Map contour level (σ) is indicated in the figure.

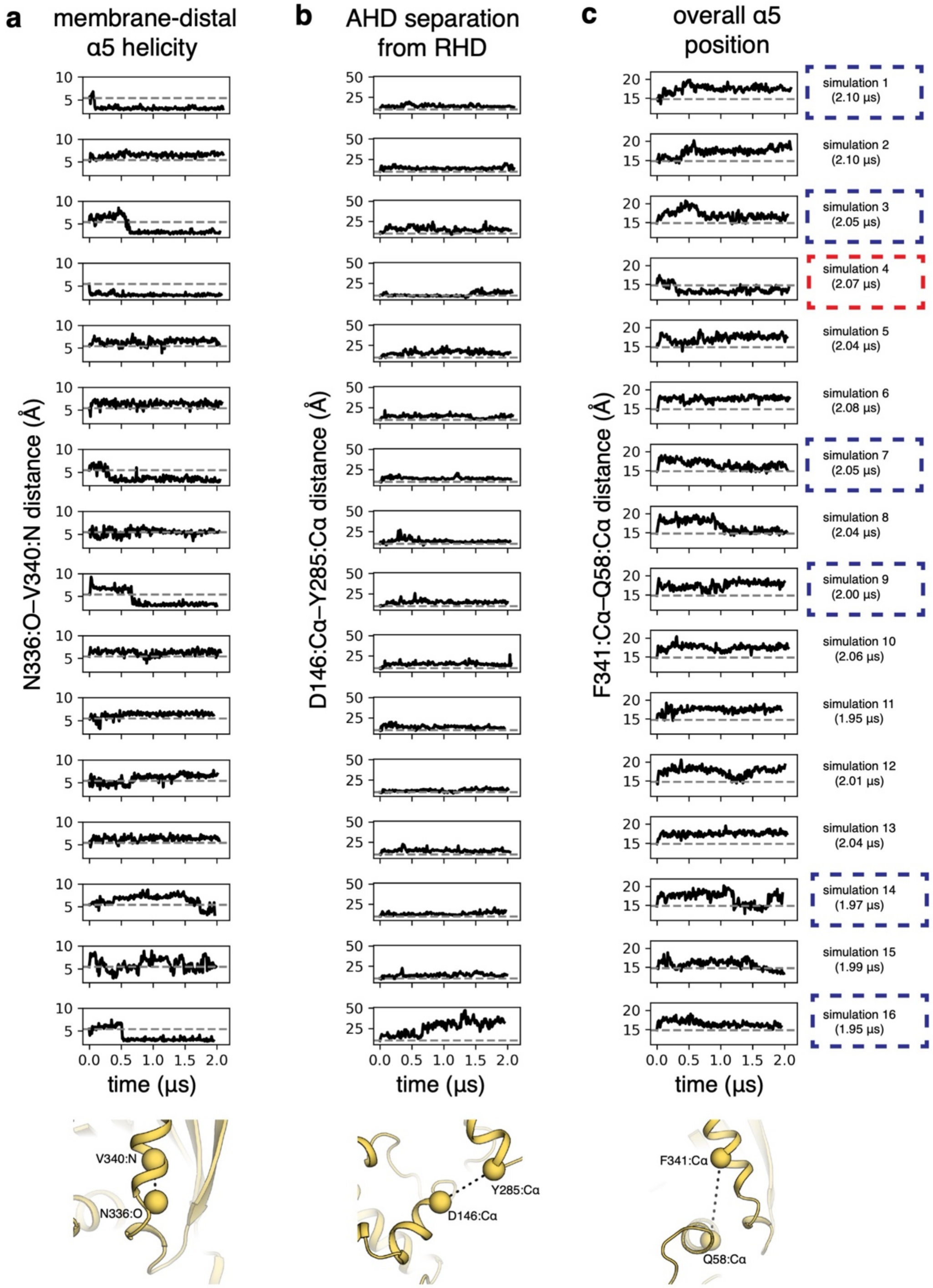


**Supplementary Fig. 6 | Metrics used to monitor conformation of Gα in MD simulations.**
These metrics, along with visual inspection, were used to identify conformational transitions. Simulations showing transitions toward the C-state are indicated by blue boxes, and the simulation showing a transition toward the E-state is indicated by a red box. **a**, Increased helicity in the membrane-distal region of the α5 helix is indicated by a decrease in the Asn336:O–Val340:N distance. **b**, AHD separation from the RHD is indicated by an increase in the Asp146:Cα–Tyr285:Cα distance. **c**, Movement of the α5 helix away from the membrane and toward GDP is indicated by a decrease in the Phe341:Cα–Gln58:Cα distance. Dashed horizontal lines represent values for the metric in the starting T2C-state structure. The renderings at the bottom of each panel show the relevant atoms in the starting structure for each metric.

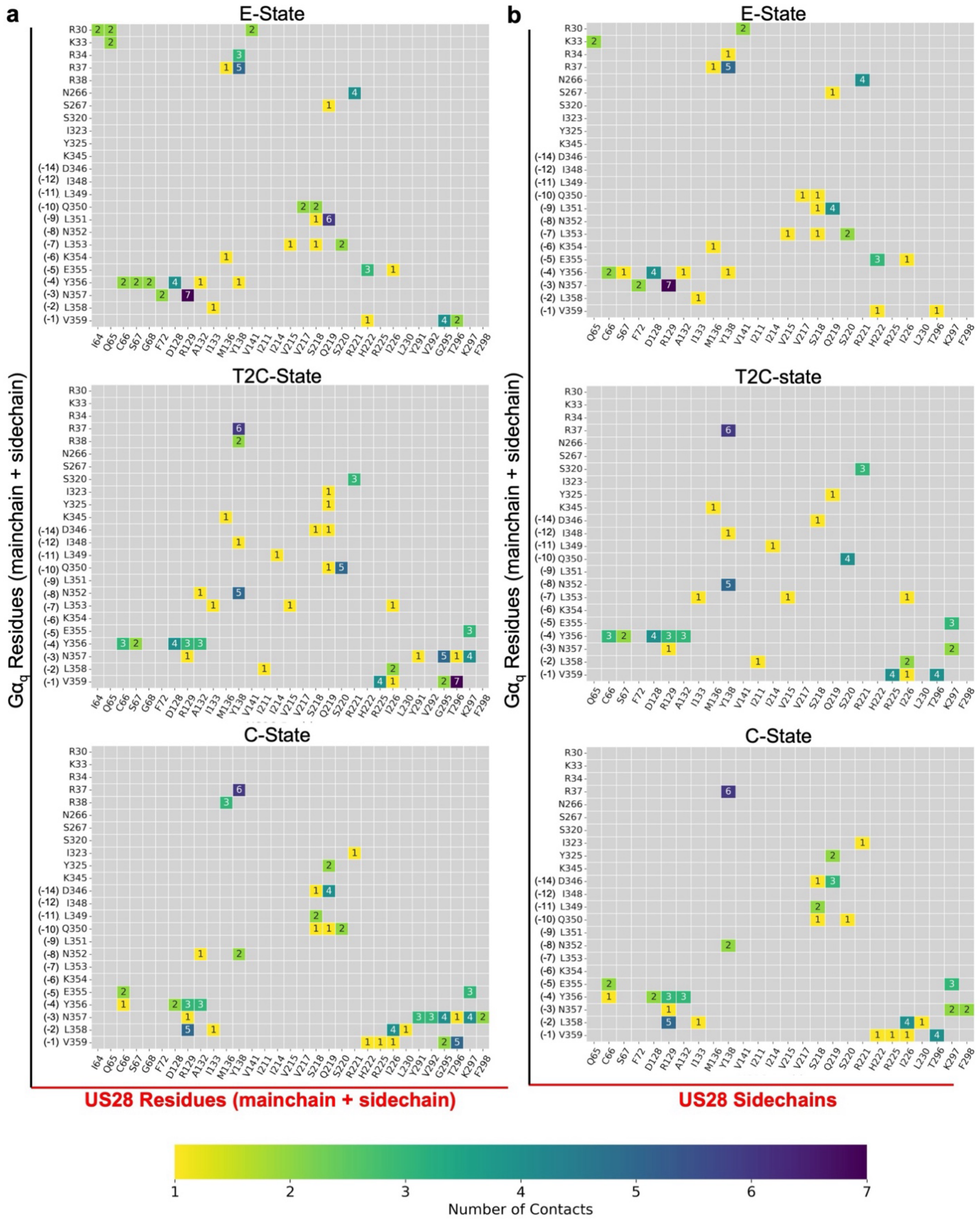


**Supplementary Fig. 7 | State-dependent interaction networks at the US28-G$_q$ interface.**
Contact maps illustrating the detailed molecular interactions between US28 residues (x-axis) and G$\alpha_q$ residues (y-axis) across the three resolved states (E-state, T2C-state, and C-state). **a**, Interactions involving both mainchain and sidechain atoms of US28. **b**, Interactions involving only the sidechain atoms of US28. In all panels, the color intensity indicates the number of atomic contacts, as shown by the scale bar.

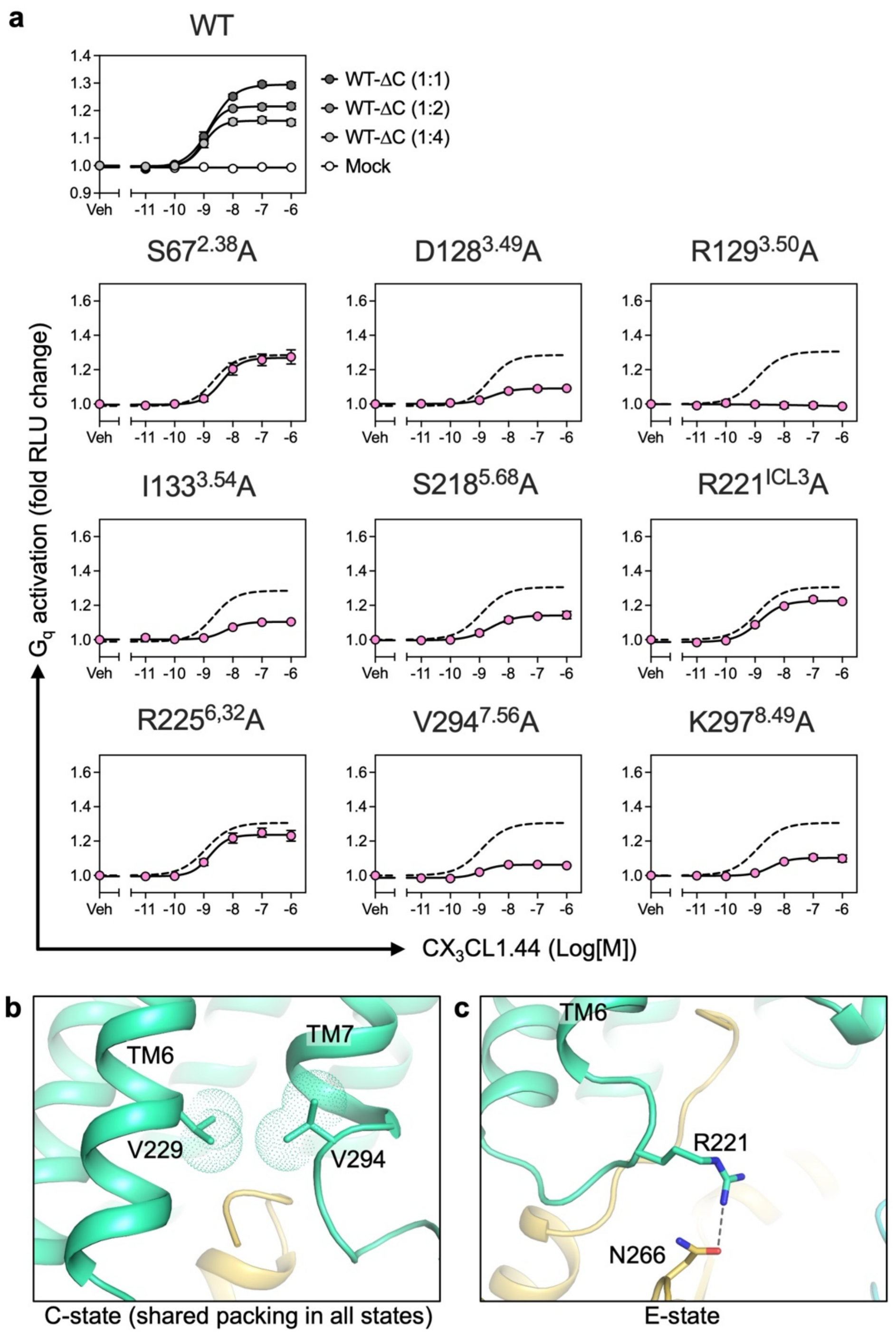


**Supplementary Fig. 8 | Dataset for the structure-guided mutagenesis analysis.**
**a**, Concentration-response curves from the NanoBiT-$G_q$ activation assay for US28Δ300 variants. Cells expressing the indicated mutants were stimulated with increasing concentrations of $CX_3CL1.44$. $G_q$ activation was monitored by the proximity signal between $G\alpha_q$-LgBiT and PLCβ2-SmBiT, measured as the fold change in relative luminescence units (RLU). Data represent the mean ± SEM from at least three independent experiments. Summarized results are shown in Fig. 4b. **b**, Val229-Val294 inter-helical packing in US28 shown in the C-state complex as an example. **c**, US28-Arg221:$G\alpha_q$-Asn266 hydrogen bond in the E-state.

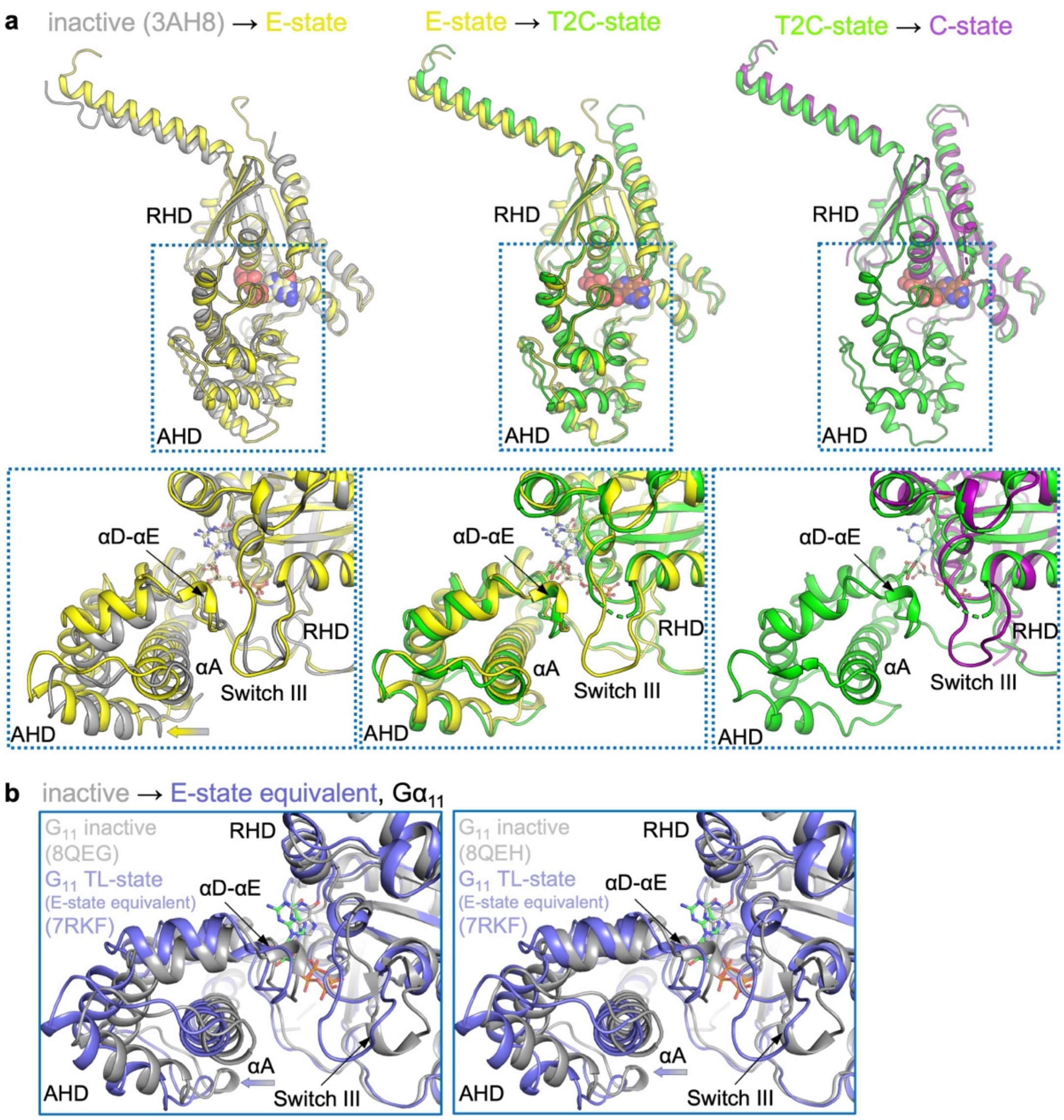


**Supplementary Fig. 9 | Overall AHD motion relative to RHD during activation.**

**a**, Structural comparison of G$\alpha_q$ showing the stepwise conformational changes leading to nucleotide release. Structures are aligned based on the RHD. (left) From the inactive to E-state, showing the slight shift for priming. (center) From the E- to T2C-states, showing the altered interdomain lock involving Switch III. (right) From the T2C- to C-states, showing complete separation of AHD. **b**, Comparisons between the inhibitor-bound inactive $G_{11}$ (gray; PDB: 8QEG, left; 8QEH, right) and $G_{11}$ in the TL-state $CX_3CL1$-US28-$G_{11}$ complex (slate blue, PDB: 7RKF, equivalent of the E-state). Both panels are oriented to provide a close-up view of the RHD-AHD interface, highlighting the repositioning of Switch III and the αA helix during the transition from the inactive to the TL-state (or E-state).

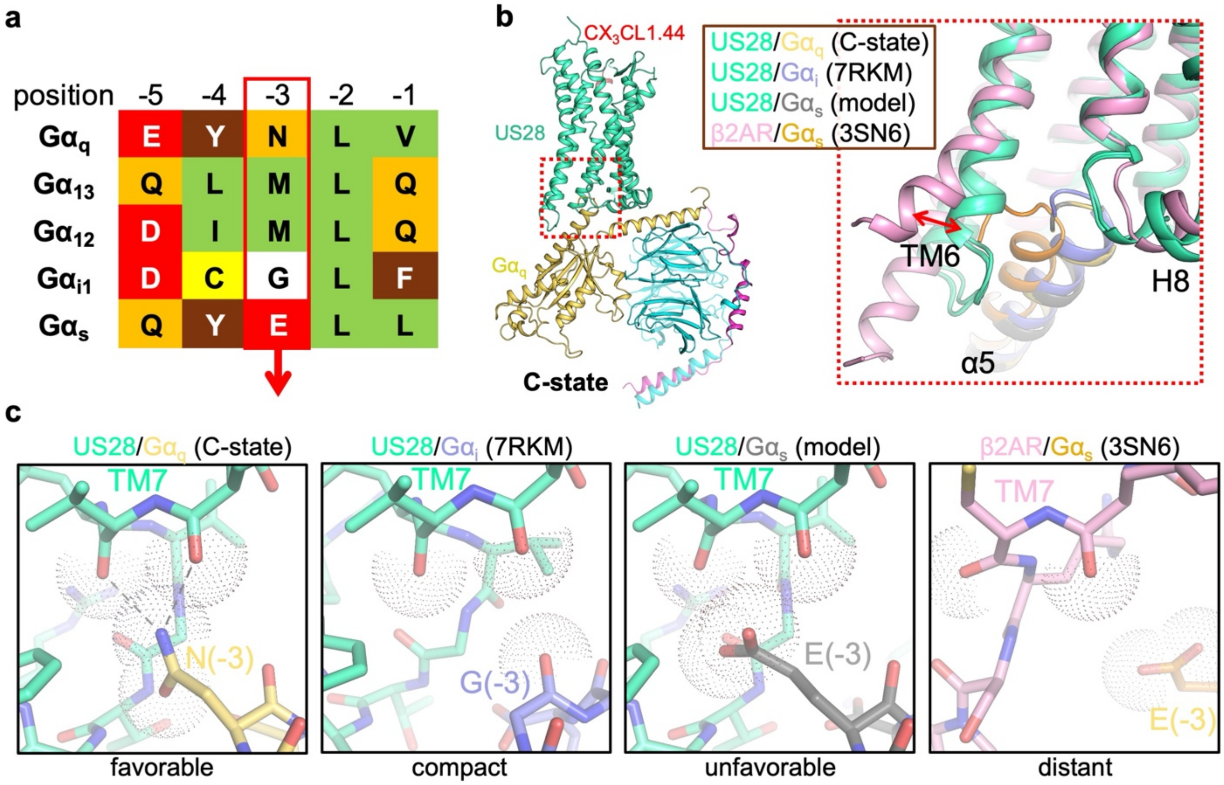


**Supplementary Fig. 10 | Structural basis for G protein subtype selectivity by US28.**

**a**, Sequence alignment of the C-terminal five residues of representative Gα subunits. Residues are color-coded by properties. $G\alpha_s$ uniquely possesses a glutamate at the -3 position. **b**, Structures of the C-state US28-$G_q$ and US28-$G_i$ complexes and a homology model of the US28-$G_s$ complex (based on the C-state US28-$G_q$ structure). **c**, Close-up views of the structures shown in panel b predict electrostatic repulsion between $G\alpha_s$-Glu(-3) and the backbone carbonyls at the intracellular end of US28 TM7, resulting from the receptor's relatively compact intracellular cavity. This contrasts with the US28-$G_q$ complex, where Asn(-3) donates hydrogen bonds to the corresponding carbonyl, and the β2AR-$G_s$ complex (PDB ID: 3SN6), where a wider opening accommodates the $G\alpha_s$ C-terminus.

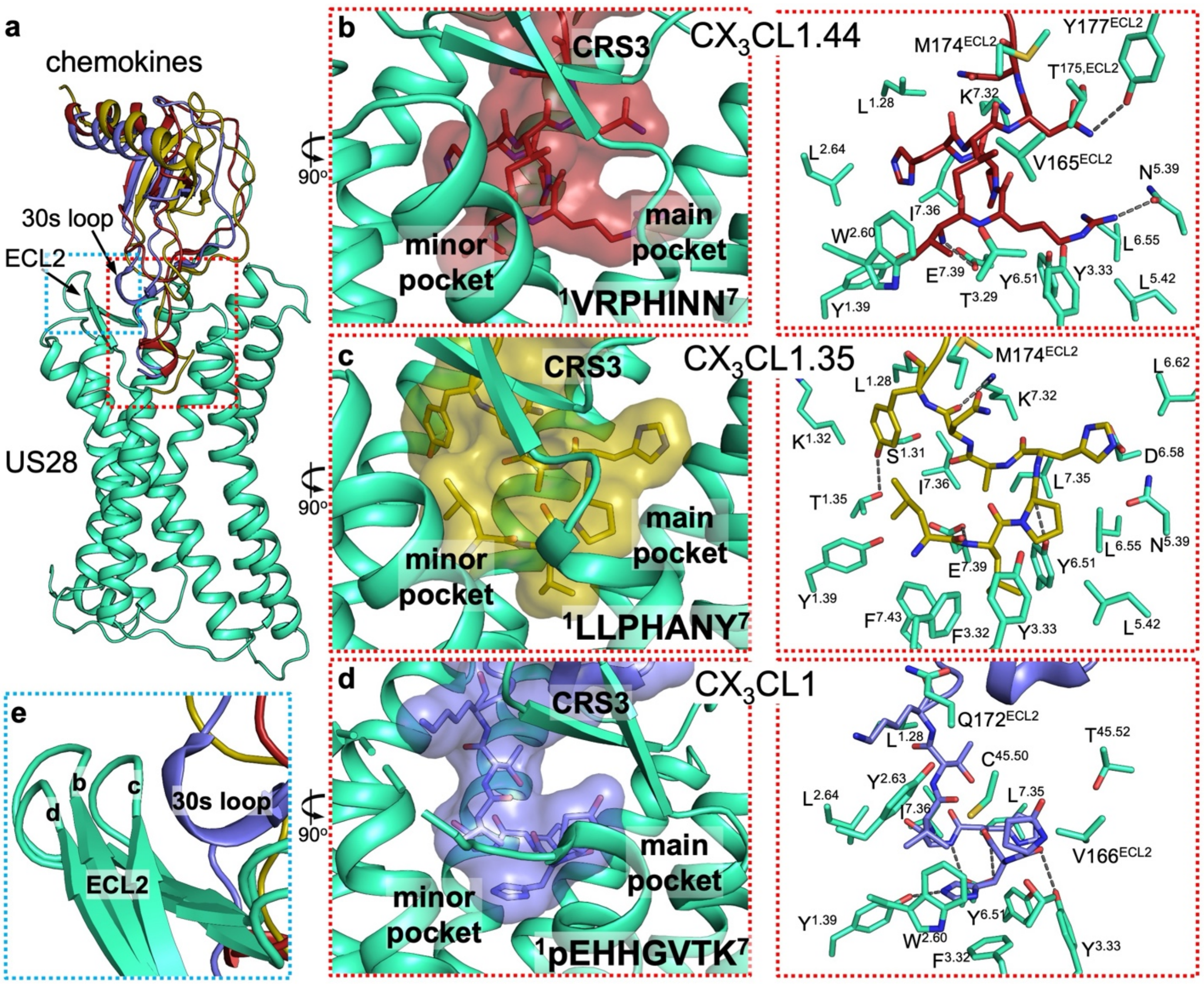


**Supplementary Fig. 11 | Ligand engagement modes.**

**a**, Overall binding modes of $CX_3CL1$ variants to US28. (**b**-**d**) Close-up views of the CRS2 interfaces between US28 and **b**, $CX_3CL1.44$ (red), **c**, $CX_3CL1.35$ (yellow), and **d**, wild-type $CX_3CL1$ (light blue). The N-terminal sequences of each chemokine are shown using one-letter codes; "pE" in panel d denotes pyroglutamate formed from the N-terminal glutamine residue. **e**, Comparison of the CRS3 interfaces between the three complexes. The labels b-d in panel e correspond to the structures shown in panels b-d.

**Supplementary References**


1. Tsutsumi, N. *et al.* Atypical structural snapshots of human cytomegalovirus GPCR interactions with host G proteins. *Sci. Adv.* **8**, eabl5442 (2022).
2. Jiao, H. *et al.* Structure basis for the modulation of CXC chemokine receptor 3 by antagonist AMG487. *Cell Discov.* **9**, 119 (2023).
3. Zhang, X. *et al.* Molecular basis for chemokine recognition and activation of XCR1. *Proc. Natl. Acad. Sci. U. S. A.* **121**, e2405732121 (2024).
4. Dror, R. O. *et al.* SIGNAL TRANSDUCTION. Structural basis for nucleotide exchange in heterotrimeric G proteins. *Science* **348**, 1361–1365 (2015).
5. Sun, X., Singh, S., Blumer, K. J. & Bowman, G. R. Simulation of spontaneous G protein activation reveals a new intermediate driving GDP unbinding. *Elife* **7**, (2018).
6. Rosenkilde, M. M., Tsutsumi, N., Knerr, J. M., Kildedal, D. F. & Garcia, K. C. Viral G protein-coupled receptors encoded by β- and γ-herpesviruses. *Annu. Rev. Virol.* **9**, 329–351 (2022).
7. Tsutsumi, N. *et al.* Insight into structural properties of viral G protein-coupled receptors and their role in the viral infection: IUPHAR Review 41. *Br. J. Pharmacol.* **182**, 26–51 (2025).
8. Waterhouse, A. *et al.* SWISS-MODEL: homology modelling of protein structures and complexes. *Nucleic Acids Res.* **46**, W296–W303 (2018).
9. Rasmussen, S. G. F. *et al.* Crystal structure of the β2 adrenergic receptor-Gs protein complex. *Nature* **477**, 549–555 (2011).
10. Flock, T. *et al.* Selectivity determinants of GPCR-G-protein binding. *Nature* **545**, 317–322 (2017).
11. Casiraghi, M. *et al.* Structure and dynamics determine G protein coupling specificity at a class A GPCR. *Sci. Adv.* **11**, eadq3971 (2025).
